# Practical observations on the use of fluorescent reporter systems in *C. difficile*

**DOI:** 10.1101/2021.06.14.448366

**Authors:** Ana M. Oliveira Paiva, Annemieke H. Friggen, Roxanne Douwes, Bert Wittekoek, Wiep Klaas Smits

## Abstract

**Objectives:** Fluorescence microscopy is a valuable tool to study a broad variety of bacterial cell components and dynamics thereof. For *Clostridioides difficile*, the fluorescent proteins CFP^opt^, mCherry^Opt^ and phiLOV2.1, and the self-labelling tags SNAP^Cd^ and HaloTag, hereafter collectively referred as fluorescent systems, have been described to explore different cellular pathways. In this study, we sought to characterize previously used fluorescent systems in *C. difficile* cells.

**Methods:** We performed single cell analyses using fluorescence microscopy of exponentially growing *C. difficile* cells harbouring different fluorescent systems, either expressing these separately in the cytosol or fused to the C-terminus of HupA, under defined conditions.

**Results:** We show that the intrinsic fluorescence of *C. difficile* cells increases during growth, independent from *sigB* or *spo0A*. However, when *C. difficile* cells are exposed to environmental oxygen autofluorescence is enhanced.

Cytosolic overexpression of the different fluorescent systems alone, using the same expression signals, showed heterogeneous expression of the fluorescent systems. High levels of mCherry^Opt^ were toxic for *C. difficile* cells limiting the applicability of this fluorophore as a transcriptional reporter. When fused to HupA, *C. difficile* histone-like protein, the fluorescent systems behaved similarly and did not affect the HupA overproduction phenotype.

**Conclusions:** The present study compares several commonly used fluorescent systems for application as transcriptional or translational reporters in microscopy and summarizes the limitations and key challenges for live-cell imaging of *C. difficile*. Due to independence of molecular oxygen and fluorescent signal, SNAP^Cd^ appears the most suitable candidate for live-cell imaging in *C. difficile* to date.

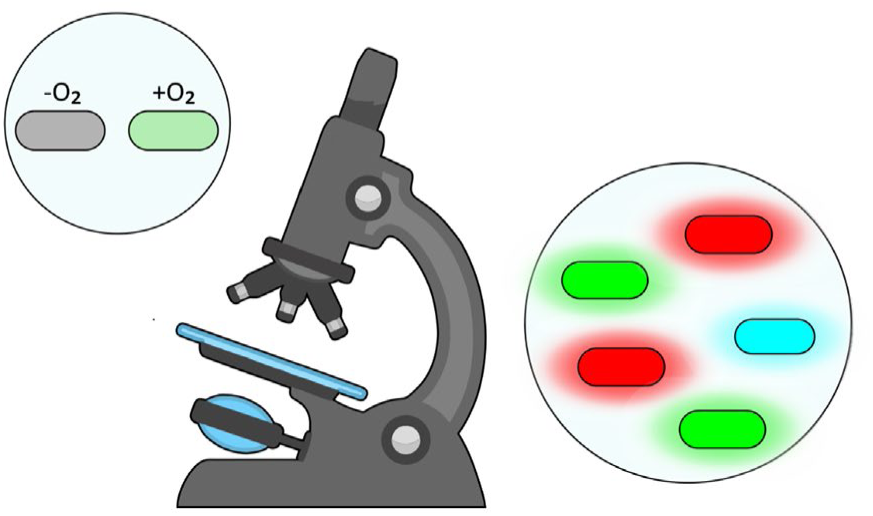

## Introduction

*Clostridioides difficile* [1] is an anaerobic gram-positive bacterium and the leading cause of antibiotic associated diarrhoea in the healthcare environment. *C. difficile* infection (CDI) occurs mainly in individuals with reduced diversity of the gut microbiota and can display a wide range of symptoms [2]. Over the past decades, the interest in the physiology of the bacterium has increased, due to the appearance of epidemic strains as well as an increase of CDI cases in the community that cannot be linked to healthcare exposure [3, 4]. *C. difficile* is a spore-forming bacterium, which allows host-to-host dissemination and enables the bacterium to persist in aerobic environments for long periods [2, 4, 5]. When the spores are ingested by the host, conditions in the small intestine trigger the spore germination and lead to the growth of the *C. difficile* vegetative cells [2, 6]. *C. difficile* vegetative cells are able to produce several factors required for the colonization of intestinal tissues and the development of the CDI, such as surface-layer proteins, which contribute to *C. difficile* adherence to the host cells, or the large clostridial toxins that disrupt the epithelial cell integrity [7, 8].

Fluorescence microscopy has been used since the early 1940‘s to detect and characterize proteins [9]. The development of a broad range of fluorescent proteins and dyes has been an important advance to a wide array of biological applications [10, 11]. Fluorescent proteins or dies have been successfully applied in the labelling of biomolecules, in gene expression and protein interaction studies, and have been used as indicators of environmental changes (e.g. pH) and as cellular stains, providing valuable insights into the dynamics of the bacterial cell [11–13]. Alongside, several microscopy techniques have been developed with increasing speed and resolution [13, 14]. Due to differences between organisms [15, 16], choosing the right fluorophore and the right imaging technique is the first step in a successful microscopy study.

In *Bacillus subtilis*, a gram-positive microorganism, the most commonly used fluorophore GFP was successfully applied to localize several proteins, such as proteins involved in replication or cell division [17]. GFP was also used to analyze gene expression, such as the expression of sigma factors important for spore development or factors that trigger biofilm formation [18, 19]. However, the use of GFP and other oxygen-dependent fluorophores is restricted to aerobic bacteria, due to the requirement of oxygen for maturation of the fluorophore [20]. This limits their applicability for studies on *C. difficile*, including live-cell imaging, as oxygen exposure imposes significant stress on anaerobic organisms, and leads to transcriptional changes that do not represent the typical growth [21–23].

Nevertheless, the use of GFP-like proteins, such as mCherry and CFP, has been shown in *C difficile* grown under anaerobic conditions. A codon-optimized version of the cyan fluorescent protein (CFP^opt^) was successfully used to localize the MldA and MldB proteins and evaluate their role on cell division [24]. The mCherry fluorophore was also codon-optimized (mCherry^Opt^) for use in *C. difficile* and was applied to localize the putative cell division protein ZapA at mid-cell [25]. It was also used as a reporter of gene expression, by placing the mCherry^Opt^ coding sequence under the control of the *pdaV* promoter (P*pdaV*), which directs transcription of the operon necessary for lysozyme resistance [25]. To allow for fluorophore maturation, fluorescence imaging was performed after prolonged exposure of the cells to environmental oxygen [24, 25].

To overcome the limitations of GFP-like molecules, oxygen-independent fluorophores have been developed that are possible candidates for use in anaerobic environments [26]. Flavin mononucleotide (FMN)-based fluorescent proteins, such as the iLOV, rely on the photoactive domain (light-oxygen-voltage) and binding of the FMN cofactor present in the bacterial cells. An iLOV version with improved photostability (phiLOV2.1) [27] has been codon-optimized for use in *C. difficile* cells to show the localization of the FtsZ protein, essential for division and septum formation, and for the FliC protein, required for cell motility [28]. Fatty-acid-binding proteins (FABPs), such as the UnaG protein, are also promising oxygen-independent fluorophores [29]. Similar to iLOV, it requires the presence of a cofactor (bilirubin) for emission of fluorescence, although in this case it is added exogenously [29]. To date, application of UnaG in *C. difficile* has not been described.

Intrinsic fluorescence of *C. difficile* cells has been previously reported with wavelengths of 500 to 550 nm, in the range of the phiLOV2.1 (green fluorescence) [25]. In *E. coli,* autofluorescence is mainly dependent on the production of small compounds, like the flavins or derivatives [30], but the source of autofluorescence in *C. difficile* is not known. However, visualization of reporters with an emission spectrum that overlaps with the autofluorescence emission wavelength can impair the effective visualization of the proteins and lead to incorrect readings in fluorescence measurements.

The need for fluorophores led to the exploration of proteins that can specifically be labelled with small chemical compounds, such as the HaloTag, SNAPtag or CLIPtag [31–33]. In *C. difficile*, simultaneous imaging of the activity of different sigma-factor dependent promoters was possible through transcriptional fusion of promoters to the codon-optimized SNAP^Cd^ and CLIP^Cd^ tags [31, 32]. The HaloTag has been used to detect and characterize the bacterial chromatin protein HupA [33].

Although the use of several fluorescent systems has been described for *C. difficile*, different experimental conditions and applications prevent a direct comparison. In this study, we benchmark technical aspects of several proteins under defined conditions, in exponentially growing *C. difficile* cells. We compare previously used fluorescent systems CFP^opt^ [24], mCherry^Opt^ [25], phiLOV2.1 [28], SNAP^Cd^ [31] and HaloTag [33] in *C. difficile*. This study highlights important limitations of existing methods and highlights key challenges for live-cell imaging of *C. difficile*.

## Materials and Methods

### Strains and growth conditions

*E. coli* strains were incubated aerobically at 37°C in Luria Bertani broth (LB, Affymetrix) supplemented with chloramphenicol at 15 µg/mL or 50 µg/mL kanamycin when required. Plasmids used in this study were maintained in *E. coli* strain DH5α. *E. coli* CA434 [34] was used for plasmid conjugation with *C. difficile* strain 630Δ*erm* [35]. Transformation of *E.coli* strains was performed using standard procedures [36].

*C. difficile* strains were incubated anaerobically in Brain Heart Infusion broth (BHI, Oxoid), with 0,5 % w/v yeast extract (Sigma-Aldrich), supplemented with 15 µg/mL thiamphenicol and *Clostridioides difficile* Selective Supplement (CDSS; Oxoid) when required. Work was performed at 37°C in a Don Whitley VA-1000 workstation or a Baker Ruskin Concept 1000 workstation with an atmosphere of 10% H_2_, 10% CO_2_ and 80% N_2_.

The growth was followed by optical density reading at 600 nm. All *C. difficile* strains used in this study are described in Table 1.

**Table 1.** *C. difficile* strains used in this study.

| Name | Relevant Genotype/Phenotype* | Origin /Reference |
| --- | --- | --- |
| wild type | 630Δ*erm*; Erm<sup>S</sup> | [35, 93] |
| IB56 | 630Δ*erm*Δ*sigB*; Erm<sup>S</sup> | [48] |
| WKS1242 | 630Δ*erm* *spo0A*::CT | [49] |
| WKS1734 | 630Δ*erm* pHEW91; Thia<sup>R</sup> | This study |
| RD14 | 630Δ*erm* pRD2; Thia<sup>R</sup> | This study |
| AF312 | 630Δ*erm* pAF308; Thia<sup>R</sup> | This study |
| RD15 | 630Δ*erm* pRD3; Thia<sup>R</sup> | This study |
| WKS1719 | 630Δ*erm* pRPF185-phiLOV; Thia<sup>R</sup> | This study |
| RD13 | 630Δ*erm* pRD1; Thia<sup>R</sup> | This study |
| AF313 | 630Δ*erm* pAF309; Thia<sup>R</sup> | This study |
| RD16 | 630Δ*erm* pRD4; Thia<sup>R</sup> | [33] |
| AF314 | 630Δ*erm* pAF310; Thia<sup>R</sup> | This study |
| RD17 | 630Δ*erm* pRD5; Thia<sup>R</sup> | This study |

\* Erm<sup>S</sup> – Erythromycin sensitive, Thia<sup>R</sup> – Thiamphenicol resistant

### Construction of the fluorescent reporter constructs

For this study, the fluorescent systems CFP^opt^, mCherry^Opt^, phiLOV2.1, HaloTag and SNAPtag^Cd^ were used. To compare the different fluorescent systems several constructs were made with only the fluorescent system or fused at the C-terminus of HupA protein (Fig. 1) [33]. All constructs were expressed under the control of the P*_tet_* promoter [37] with the *cwp2* ribosomal binding site (aaatttgaattttttagggggaaaatac). For the C-terminal fusions to the HupA protein, a short GS linker (SGSGSGS) was introduced, as previously described [33]. All the vectors and oligonucleotides used in this study are listed in Tables 2 and 3, respectively. All the constructs were confirmed by Sanger sequencing.

**Fig. 1.**
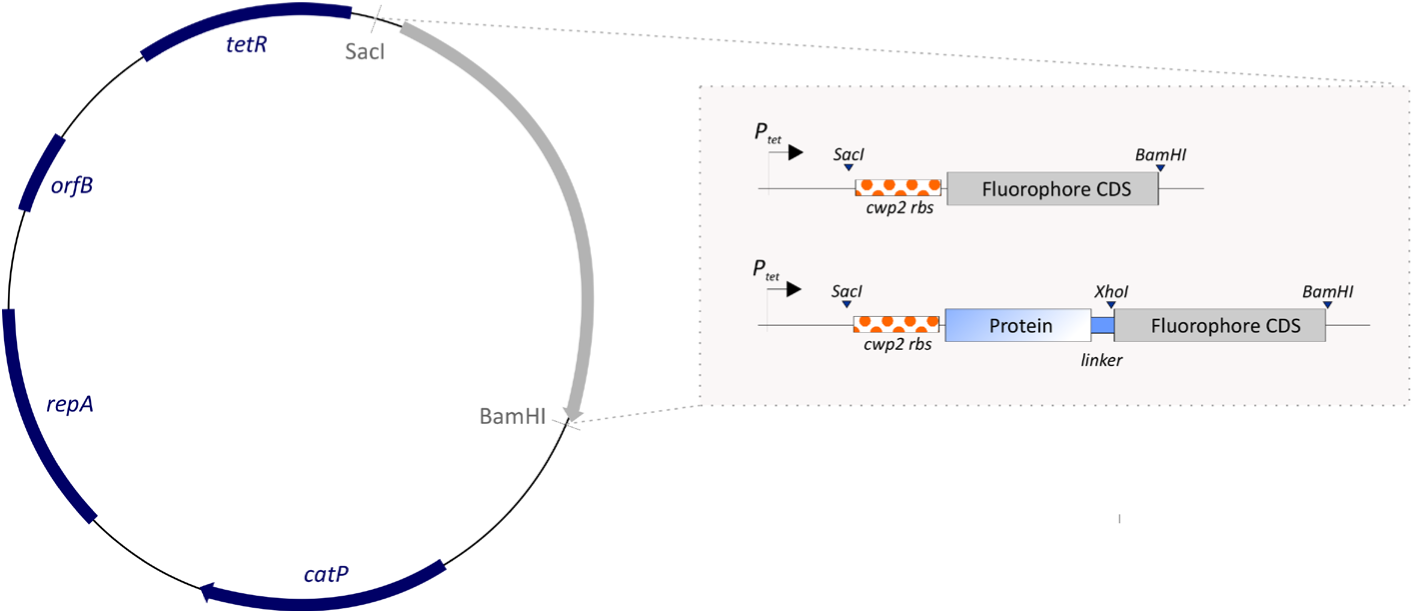
Representation of the modular vector for the expression of different fluorescent systems. The pRPF185 backbone vector used is depicted. Genes present are denoted in dark blue (*catP, repA, orfB, tetR*) [37]. The modular cassette (grey arrow), under the control of the P*_tet_* promoter, for both the expression of the fluorescent systems cytosolic (grey boxes) or for fusions at the N-terminus to the study protein (blue-white box). The positions of used restriction sites are marked (*ScaI*, *XhoI* and *BamHI*). The *cwp2* ribosomal binding site (orange dotted box) and the GS linker (blue box) are depicted.

**Table 2.**
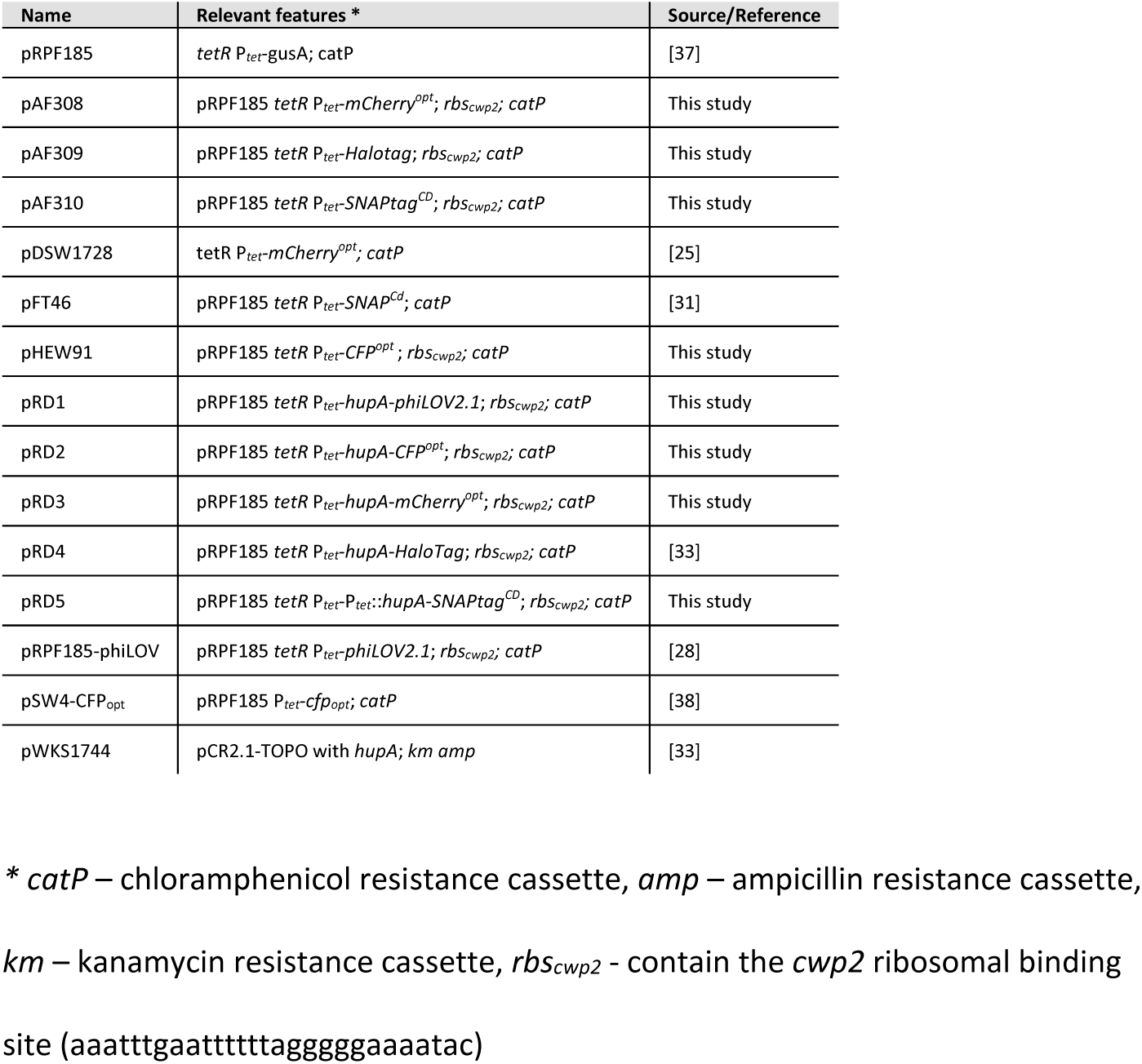
Vectors used in this study, alphabetically.

**Table 3.**
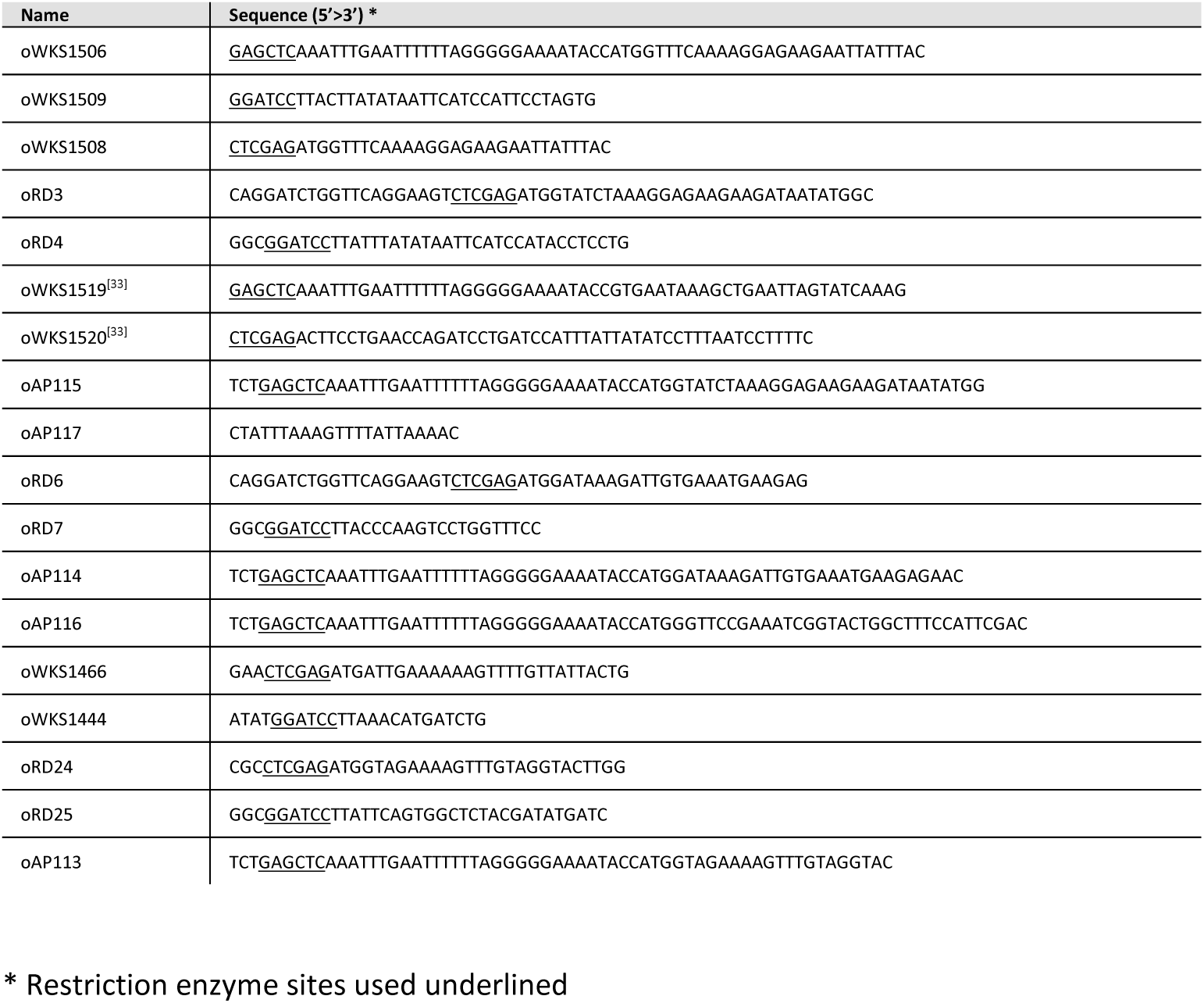
Oligonucleotides used in this study.

For the CFP^opt^ constructs, the *cfp* gene was amplified from pSW4-CFP_opt_ [38], obtained from the Bacillus Genetic Stock Center (http://www.bgsc.org/), with the primer set oWKS1506/oWKS1509. The amplified fragment was digested with *SacI* and *BamHI*, yielding vector pHEW91, for cytosolic CFP^opt^ expression. To create the HupA fusion to the CFP^opt^ protein, the CFP^opt^ coding sequence was amplified with the primer set oWKS1508/oWKS1509. The amplified fragment was digested with the restriction enzymes *XhoI* and *BamHI* and ligated into the pRD4 vector [33] digested with the same enzymes, yielding vector pRD2, encoding for HupA-CFP^opt^.

To create the cytosolic version, the gene encoding for *mCherry^Opt^* from pDSW1728 [25], kindly provided by Dr. Eric Ransom, was amplified with the primer set oAP115/oAP117. The product was digested with *SacI* and *BamHI* and ligated into pRPF185 digested with the same restriction enzymes, yielding plasmid pAF308. To generate the fusion to mCherry^Opt^ protein, the gene encoding for *mCherry^Opt^* was amplified with the primer set oRD3/oRD4 from pDSW1728. The *hupA* gene was amplified from pWKS1744 with the primer set oWKS1519 and oWKS1520. Overlap extension PCR was used to create *hupA-mCherry^Opt^*, according to manufacturer instructions (Accuzyme) with primer set oWKS1519/oRD4. The resulted fusion PCR product was digested with *SacI* and *BamHI* and ligated into pRPF185 digested with the same restriction enzymes, yielding the plasmid pRD3.

The cytosolic phiLOV2.1 is encoded on the pRPF185-phiLOV vector [28], kindly provided by Dr. Gillian Douce. To create the HupA-phiLOV2.1 expression vector, the gene encoding for the codon-optimized phiLOV2.1 was amplified from the pRPF185-phiLOV vector with the primer set oWKS1466/oWKS1444. The amplified fragment was digested with *XhoI* and *BamHI* and inserted onto pRD4, digested with the same enzymes yielding vector pRD1.

For the cytosolic HaloTag expression, the *haloTag* coding sequence was amplified from pRD4 with primer set oAP116/oAP117. The amplified fragment was digested with *SacI* and *BamHI* and ligated to the pRPF185 digested with same enzymes, yielding vector pAF309. HupA-HaloTag expression is encoded in pRD4, previously used [33].

To create the cytosolic version, the gene encoding for *SNAP^Cd^* was amplified from pFT46 [31], kindly provided by Dr. Adriano Henriques, with the primer set oAP114/oAP117. The product was digested with *SacI* and *BamHI* and ligated into pRPF185 digested with the same restriction enzymes, yielding plasmid pAF310. For the *hupA-SNAP^Cd^* construct, the gene encoding for *SNAP^Cd^* was obtained with the primers oRD6 and oRD7 from pFT46. The *hupA* gene was amplified from pWKS1744 with the primer set oWKS1519/oWKS1520. The overlap extension PCR was performed using the primer set oWKS1519/oRD7, according to manufacturer instructions (Accuzyme). The resulted fusion PCR product was digested with *SacI* and *BamHI* and ligated into pRPF185 digested with the same restriction enzymes, yielding plasmid pRD5.

### Fluorescent microscopy

*C. difficile* strains were cultured in BHI/YE, with thiamphenicol when required. When necessary cells were induced with 200 ng/mL ATc for 1 hour at an OD_600nm_ of approximately 0.4. The sample preparation for fluorescence microscopy was carried out under anaerobic conditions.

For the fluorescence microscopy, 1 mL culture was collected and centrifuged (2000×g, 1min). Cells were re-suspended in 1 mL pre-reduced PBS and centrifuged (2000g, 1min). When required, cells were incubated with 100 nM Oregon Green substrate for HaloTag (G280A, Promega) or 100 nM TMR substrate for SNAPtag (S9105S, NEB), for 30 min anaerobically. For oxygen-dependent fluorescent systems, samples were exposed aerobically for 30 min. After incubation cells were centrifuged (2000g, 1min). When required samples were left in PBS for 15 or 30 min anaerobically or exposed to oxygen, when necessary, until imaging. 900 µL supernatant was removed and cells re-suspended in the leftover supernatant. When necessary samples were incubated with 1 μM DAPI (Roth), for 1min and centrifuged (2000×g, 1min). 50 µL supernatant was removed and cells re-suspended in the leftover supernatant. 2 µL sample were spotted on 1.5% agarose patches and imaged within 2 to 15 min. 2 µL ProLong Gold Antifade Mountant (S36936, Invitrogen) was added on top the sample, for HaloTag imaging when required.

Samples were imaged with a Leica DM6000 DM6B fluorescence microscope (Leica) equipped with DFC9000 GT sCMOS camera using an HC PLAN APO 100x/1.4 OIL PH3 objective, using the LAS X software (Leica). The filter sets used for imaging are summarized in Table 4. Images were acquired with an exposure time of 200 ms for all the channels used and an intensity value of 157 for phase contrast and 5 for the fluorescent filters.

**Table 4.** Filter sets and correspondent fluorescent dyes/ fluorescent systems used in this study.

| Filter set<br>(Leica<br>no.) | Excitation filter | Dichroic<br>mirror | Emission filter | Fluorescent system /dye | Ex<br>(max) | Em<br>(max) |
| --- | --- | --- | --- | --- | --- | --- |
| DAPI<br>11504203 | BP 350/50 | LP 400 | BP 460/50 | DAPI | 357 | 457 |
| CFP<br>11504163 | 436/20 | 455 | 480-40 | CFP<sup>opt</sup> | 435 | 475 |
| Alexa<br>488/L5<br>11504166 | 480/40 | 505 | 527/30 | phiLOV2.1 | 450 | 492 |
|  |  |  |  | Halotag - Oregon Green | 496 | 516 |
| Y3<br>11504169 | 545/25 | 565 | 605/70 | SNAP<sup>Cd</sup> - TMR | 554 | 580 |
|  |  |  |  | mCherry<sup>Opt</sup> | 585 | 610 |

### Data analysis

TIFF exported pictures with raw data where exported for analysis from LAS X software. No picture treatment was performed for image analysis.

Data was analysed with MicrobeJ package version 5.13I [39] with ImageJ 1.52p software (https://imagej.nih.gov/ij/index.html) [40]. Recognition of cells was limited to specific settings: 2 - 16 µm^2^ area, 0.4 - max µm range of width, 0 - 0.7 circularity, 0 - 0.16 curvature and 0 - 0.3 angularity. Cells with defective detection were excluded from analysis.

Analysis of autofluorescence was through the intensity profile analysis of the mean intensity throughout the longitudinal axis of the bacterial cell in the green channel (Alexa 488/L5 filter, Table 4).

Fluorescence associated with the fluorescent system and nucleoid staining where detected with stain feature separately, with a Z-score of 3 and 5, respectively. Cells with no nucleoid staining were excluded from analysis. Raw data was exported and visualized with Prism 8.3.1 (GraphPad, Inc, La Jolla, CA). Statistical analysis was performed in Prism 8.3.1 (GraphPad, Inc, La Jolla, CA), with the Kruskal-Wallis statistical test for analysis of samples with a variance not normally distributed, with p<0.0001.

Representative pictures where selected and cropped in LAS X software (Leica). Contrast increase was performed equally for better visualization.

### In-gel fluorescence

*C. difficile* strains were cultured in BHI/YE, and when appropriate induced at an OD_600nm_ of 0.3-0.4 with 200 ng/mL ATc concentration for up to 1 hour. Samples were collected and centrifuged at 4°C. Pellets were resuspended in 100 µL Lysis Buffer (10 mM Tris-HCl pH 7.5, 10 mM EDTA, protease cocktail inhibitor (Roche), 0.1 mg/mL lysozyme) and incubated for 30 min at 37°C. Samples were incubated with 100 nM OregonGreen substrate for HaloTag (Promega) or 100 nM TMR substrate for SNAPtag (NEB) for 30 minutes at 37°C, when necessary. Loading buffer (250 mM Tris-Cl pH 6.8, 10 % SDS, 10% β-mercaptoethanol, 50% glycerol, 0.1% bromophenol blue) was added to the samples without boiling and samples were run on 12% SDS-PAGE gels. Gels were imaged with Uvitec Alliance Q9 Advanced machine (Uvitec) with the filters F-535 (460 nm), F-595 (520 nm) and F-695 (650nm).

### Spot assay

Cells were grown until OD_600_ of 1.0 in BHI/YE with thiamphenicol when required. The cultures were serially diluted (100 to 10^−5^) and 2 µL from each dilution were spotted on pre-reduced BHI/YE plates supplemented with CDSS, thiamphenicol and 200 ng/mL ATc when appropriate. Plates were imaged after 24 hours anaerobic incubation at 37°C.

All results were combined for publication with Prism 8.3.1 (GraphPad, Inc, La Jolla, CA) and CorelDRAW X8 (Corel).

## Results

### *C. difficile* autofluorescence increases in stationary phase

To better understand the intrinsic fluorescence of *C. difficile,* we monitored the optical density and the fluorescence of a wild type *C. difficile* 630Δ*erm* culture for 24 hours. Cells were grown in a rich medium (BHI/YE), from a pre-culture and samples were taken at different times during cell growth (Fig. 2A). To assess the intrinsic fluorescence, samples were collected and immobilized on an agarose patch under anaerobic conditions. Microscopy of cells in the middle of the patch was performed in aerobic conditions within 5 minutes after removal from the anaerobic cabinet in order to limit oxygen influence and damage of the cells. Handling of the samples was kept to a minimum and thus no dye was added for membrane or nucleoid staining.

**Fig. 2.**
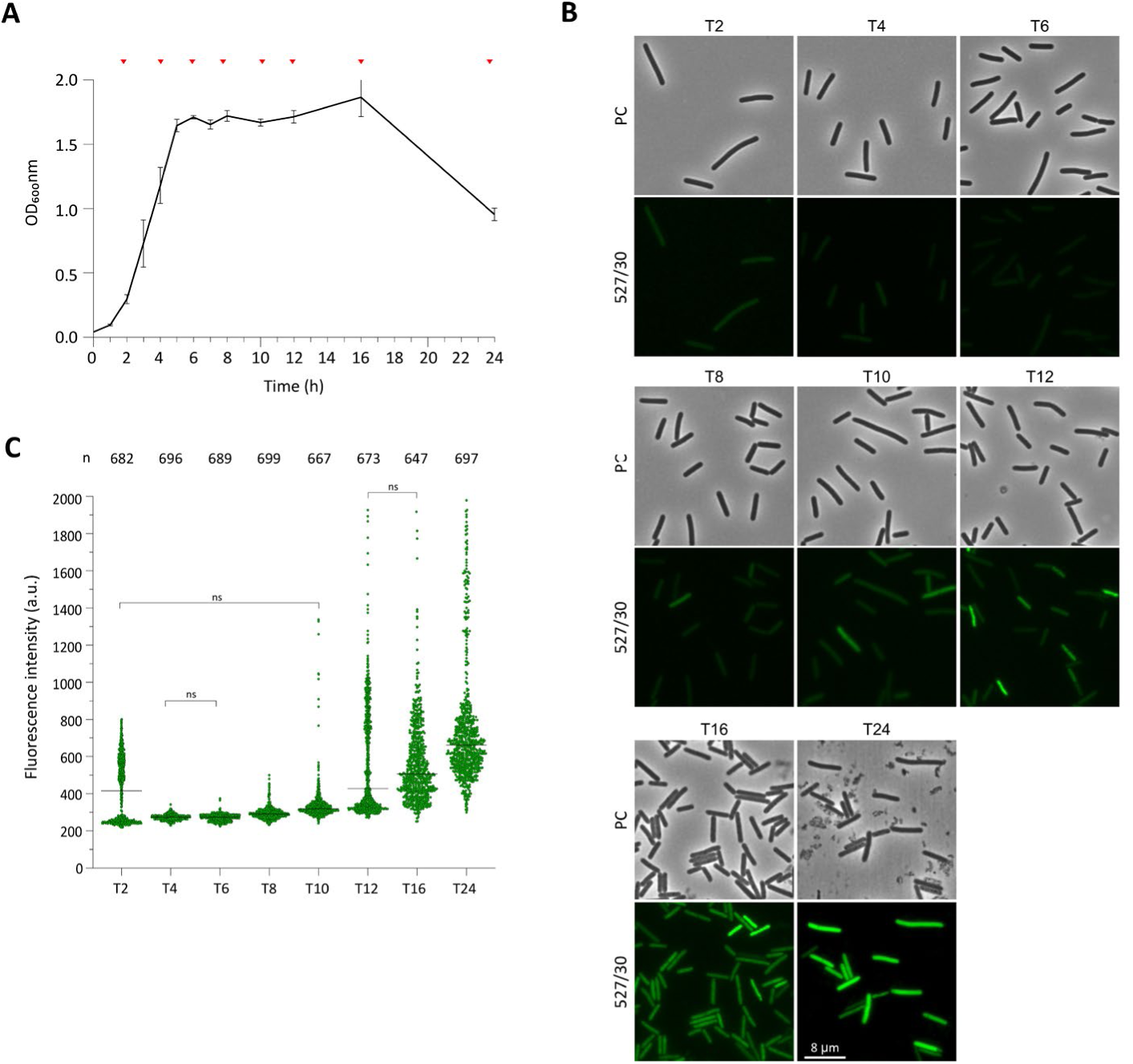
*C. difficile* autofluorescence increases over time. **A)** Growth curve of *C. difficile* strain 630Δ*erm*, incubated anaerobically at 37 °C and followed for 24 hours. Optical density was measured at a wavelength of 600 nm (OD_600nm_). Time points analyzed are indicated by red arrows. **B)** Fluorescence microscopy analysis of *C. difficile 630Δerm* at the indicated time points (2, 4, 6, 8, 10, 12, 16 and 24 hours post-inoculation). The effect was verified by three independent microscopy experiments performed on different days. Phase contrast (PC) and green channel (emission filter 527/30 nm, for autofluorescence) are shown. Because growth is asynchronous in these conditions, cells representing different cell cycles stages can be observed. Scale bar = 8 µm. **C)** Dot plots of mean longitudinal green fluorescence (a.u.) of the analysed *C. difficile* 630Δ*erm* cells. Black lines represent the median values. Quantifications were performed using MicrobeJ from at least two biological replicates for each condition. The number of cells analysed per time point (n) is indicated above each graph. All results were statistically significant with p<0.01 by one-way ANOVA, except when indicated otherwise (ns).

During exponential phase, no considerable fluorescence signal in the green channel (emission wavelength 527/30) was detected (<300 A.U., Fig. 2B - C, T4 to T8). In contrast, we observed a significant increase in the fluorescence signal in stationary growth phase (>300 A.U., Fig. 2B - C, T10 to T24), reaching higher fluorescent intensity signals at 24 hours (758.9 ± 347.5 A.U., Fig. 2C, T24). These results reconcile contrasting observations from previous studies, where actively dividing cells exhibit low levels of green fluorescence [33], but high fluorescent signals were reported for stationary phase cells [31].

Quantification of the images shows a 2-3-fold increase in the average fluorescence intensity in stationary growth phase, with a large heterogeneity between cells (Fig. 2B - C). Growth is non-synchronous for *C. difficile* and in stationary phase both nutrient depletion and developmental reprogramming occurs, which could contribute to the observed heterogeneity.

We also observed a broader range of signal intensities at 2 hours (416.1 ± 167.4 A.U., Fig. 2B and C, T2) which we attribute to carry-over from the 12-hour preculture, as it disappears as cells start to grow exponentially and with a similar variability as in the stationary phase.

### Autofluorescence is independent of *sigB* and *spo0A*

Growth dependent autofluorescence has also been observed for organisms such as *E. coli* or *Bacillus pumilus* [30, 41]. In *E.coli,* riboflavin and riboflavin-derivative compounds are the main components responsible for the green fluorescence [30]. Riboflavin is an essential precursor of compounds such as the flavin mononucleotide (FMN) and flavin adenine dinucleotide (FAD) that are cofactors of flavoproteins. Flavoproteins have pleiotropic functions [42] and have been implicated in the oxidative stress response in several anaerobes [43, 44]. In *C. difficile,* flavoproteins are overrepresented in the regulon of the general stress response sigma factor σB [45, 46] and have been implicated in spore resistance [47].

To determine whether regulators involved in oxidative stress and sporulation contribute to the increase in autofluorescence in stationary growth phase, two mutants lacking the σ^B^ gene (Δ*sigB*) or carrying a disrupted *spo0A* gene (*spo0A*::CT) were used [48, 49]. σ^B^ is the alternative sigma factor that directs the general stress response in several bacteria [50] and is implicated in metabolism, sporulation and stress response in *C. difficile* [45, 46]. Spo0A is a highly conserved transcriptional regulator that controls the onset of sporulation but also regulates other virulence associated factors such as motility and metabolism [49, 51].

Both mutants were grown under the same conditions as the wild type *C. difficile* strain 630Δ*erm*, previously mentioned, and no differences in growth were observed between the two mutants (Fig. 3A), or in comparison with the wild-type (Fig. 2A).

**Fig. 3.**
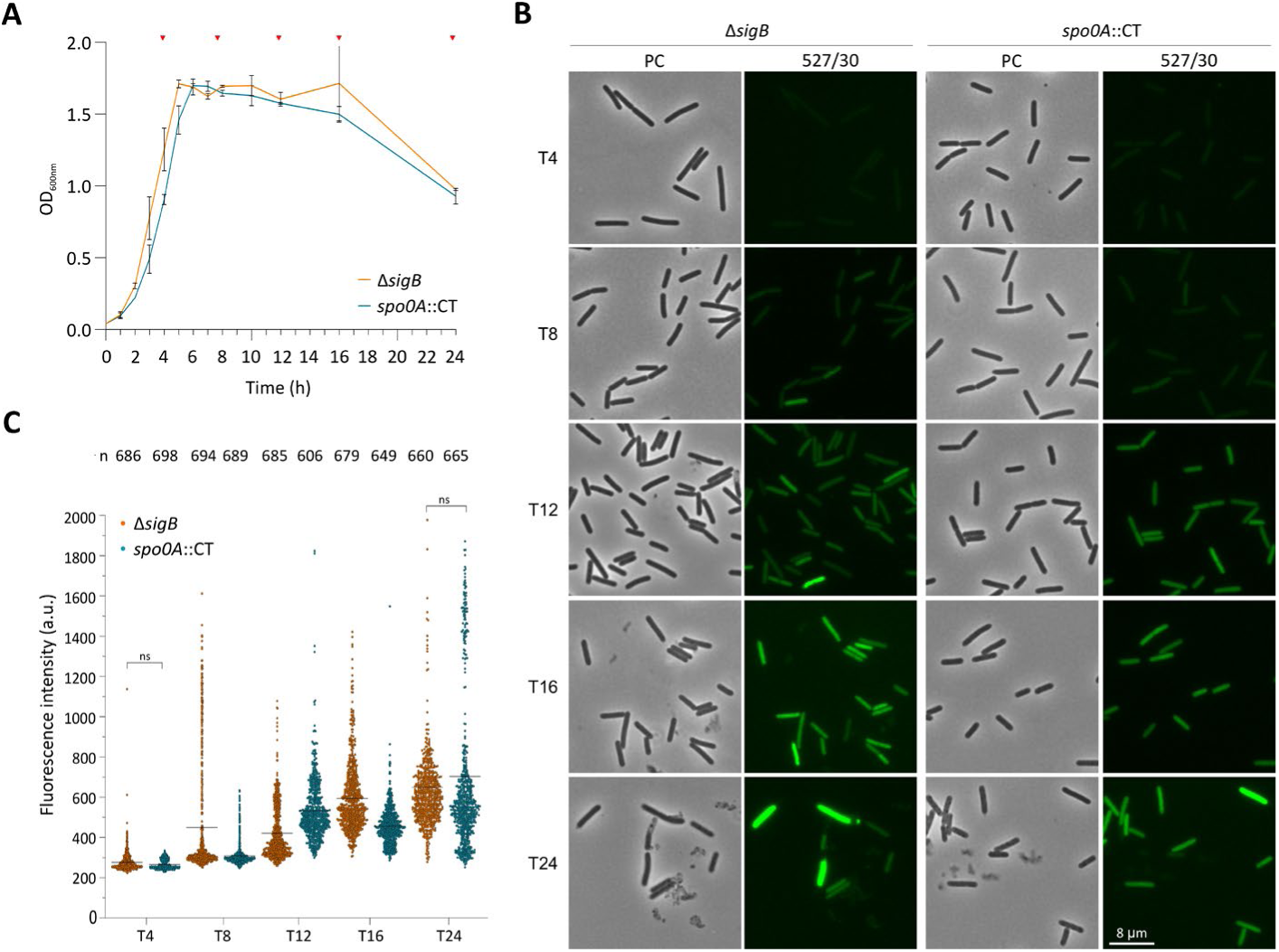
Autofluorescence in the *sigB* and *spo0A* mutants. **A)** Growth curve of *C. difficile* strains IB56 (Δ*sigB*, orange line) and WKS1242 (*spo0A*::CT, blue line) incubated anaerobically at 37 °C and followed for 24 hours with measurement of optical density at 600 nm (OD_600nm_). Time points analyzed are indicated with a red arrow. **B)** Fluorescence microscopy analysis of *C. difficile* strains at the indicated time points (4, 8, 12, 16 and 24 hours). The effect was verified by three independent microscopy experiments. Phase contrast (PC) and green channel (emission filter 527/30 nm, for autofluorescence) are shown. Because growth is asynchronous in these conditions, cells representing different cell cycles stages can be observed. Scale bar = 8 µm. **C)** Dot plots of mean longitudinal green fluorescence (a.u.) of the analysed *C. difficile* cells Δ*sigB* (orange) and *spo0A*::CT (blue). Black lines represent the median values. Quantifications were performed using MicrobeJ from at least two biological replicates for each condition. The number of cells analysed per time point (n) is indicated above each graph. All results were statistically significant with p<0.01 by one-way ANOVA, except when indicated otherwise (ns).

Similar to wild type, a negligible fluorescence signal was detected in exponentially growing cells (T4 to T8, Fig. 3B - C, Fig. S1), whereas both the strength of the signal and the variability increased significantly at later timepoints (T12 to T24, Fig. 3B - C, Fig. S1).

Taken together, these results suggest that *sigB* and *spo0A* do not have a major influence on the autofluorescence of *C. difficile*. Though we cannot exclude a contribution of (ribo)flavin, these results suggest that, if such a mechanism exists, it is unlikely to be mediated through transcriptional regulation by the general stress response sigma factor or sporulation-specific regulators that act downstream of Spo0A.

### Autofluorescence increases in the presence of oxygen

Autofluorescence uncoupled from the sporulation process has also been reported for *B. pumilus*. In this organism, the production of a pigment responsible for autofluorescence was found to be influenced by environmental cues such as growth conditions [41]. Interestingly, the pigment is also involved in resistance to H_2_O_2_ [41], suggesting a link between autofluorescence and oxidative stress.

To better evaluate the possible contribution of oxygen to *C. difficile* autofluorescence, we took samples in exponential growth phase with an approximate optical density (OD_600nm_) of 0.45 and imaged them anaerobically, or after exposure to oxygen. Anaerobically incubated samples demonstrated a low mean fluorescence intensity (322.7 ± 52.75 A.U., Fig. 4, -O_2_), with a distribution in line with our previous experiments (Fig. 2, T2 and T4). However, when cells were exposed for 15 min to environmental oxygen a significant increase in fluorescence was observed (801.9 ± 134.8 A.U., Fig. 4, +O_2_). Notably, the levels of autofluorescence upon oxygen exposure are equal or higher than those observed in stationary phase cultures of *C. difficile*. These results suggest that the presence of oxygen can significantly contribute to the observed autofluorescence of *C. difficile*.

**Fig. 4.**
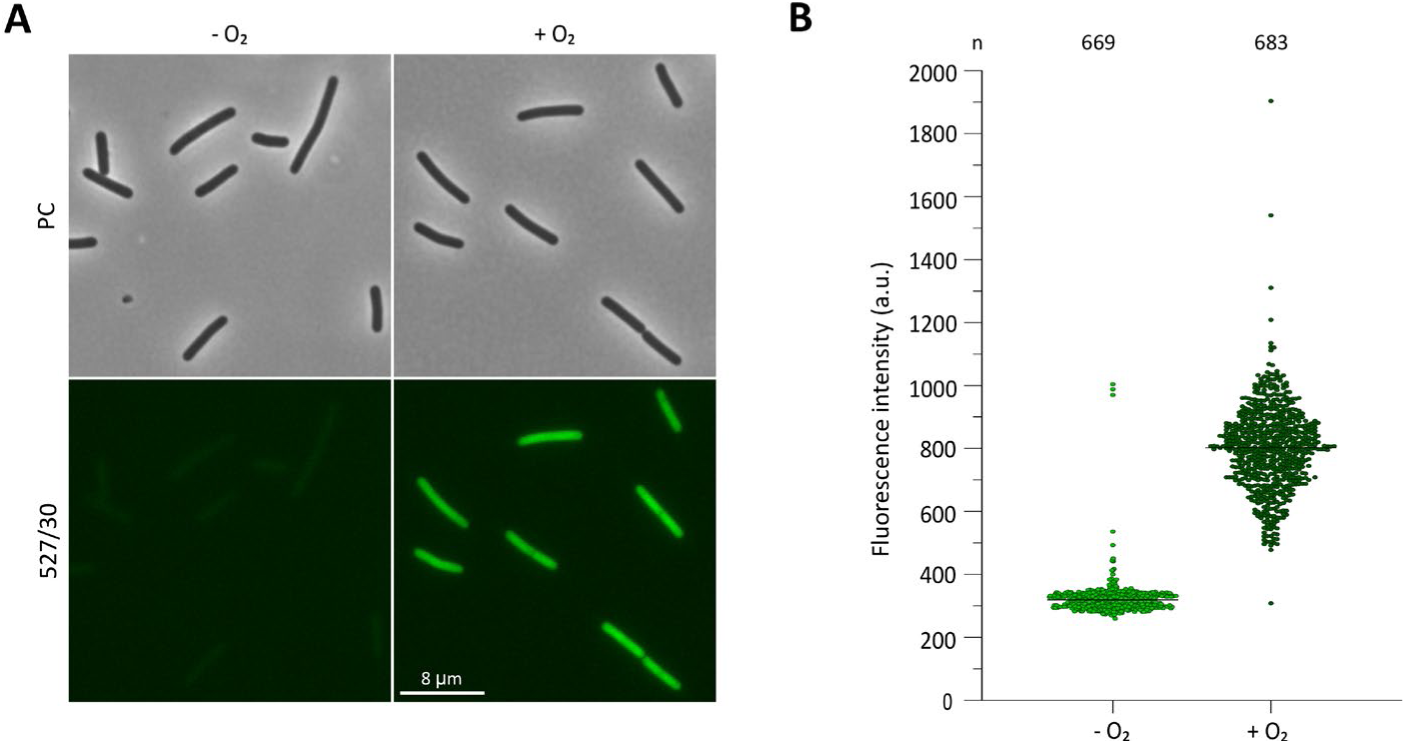
Autofluorescence is dependent on oxygen exposure. **A)** Fluorescence microscopy analysis of *C. difficile* strains incubated for 15 min in aerobic (+O_2_) or anaerobic conditions (-O_2_). The effect was observed in three independent microscopy experiments. Phase contrast (PC) and green channel (emission filter 527/30 nm, for autofluorescence) are shown. Because growth is asynchronous in these conditions, cells representing different cell cycles stages can be found. Scale bar = 8 µm. **B)** Dot plots of mean longitudinal green fluorescence (a.u.) of the analysed *C. difficile* cells -O_2_ (light green) and +O_2_ (dark green). Black lines represent the median values. Quantifications were performed using MicrobeJ from at least two biological replicates for each condition. The number of cells analysed per time point (n) is indicated above each graph. The difference is statistically significant with p<0.01 by one-way ANOVA.

### Overexpression of mCherry^Opt^, but not other fluorescent systems, in the cytosol is toxic to *C. difficile*

Having established conditions for imaging *C. difficile* with low background fluorescence, we continued to evaluated the different commonly used fluorescent systems CFP^opt^, mCherry^Opt^, phiLOV2.1, HaloTag and SNAP^Cd^ [24, 25, 28, 31, 33]. To enable comparison of fluorescent systems, we constructed modular vectors expressing the selected fluorescent systems, separately (i.e. cytosolically expressed, Fig. 1). To exclude a confounding effect of differing expression signals, all constructs contained the same inducible promoter (P*_tet_*) with identical ribosome binding sites (derived from the *cpw2* gene) [37].

To evaluate the effect of overexpression of the fluorescent systems on the viability of the *C. difficile* cells, we performed a spot-assay. The different *C. difficile* strains were spotted in 10-fold serial dilutions on BHI/YE agar with *C. difficile* selective supplement (CDSS), supplemented with thiamphenicol and anhydrotetracycline (ATc) when appropriate. All the strains grew indistinguishably until a dilution of 10^−5^ on medium lacking thiamphenicol and ATc (Fig. 5, top panel). As expected, the wild type strain (630Δ*erm*, wt) is unable to grow in the presence of thiamphenicol (Fig. 5, middle panel), but all other strains are. When grown in the presence of both thiamphenicol and 200 ng/ml ATc, striking differences are observed between the strains (Fig. 5, lower panel). Viability upon overexpression of CFP^opt^, phiLOV2.1 and SNAP^Cd^ does not markedly differ from that under non-inducing conditions. A minor effect on viability is observed when HaloTag is overexpressed, with visible growth until dilution 10^−4^. However, overexpression of mCherry^Opt^ shows a drastic effect on the viability with a 4-log reduction compared to the other fluorescent systems. This effect was not previously reported [25]. It is conceivable that the use of the strong *cwp2* ribosomal binding site here leads to higher expression compared to previous work [25]. Nonetheless, our work highlights that for use as a cytosolic reporter (for instance for gene expression), one should consider possible negative effects of reporter expression, at least in the case of mCherry^Opt^.

**Fig. 1.**
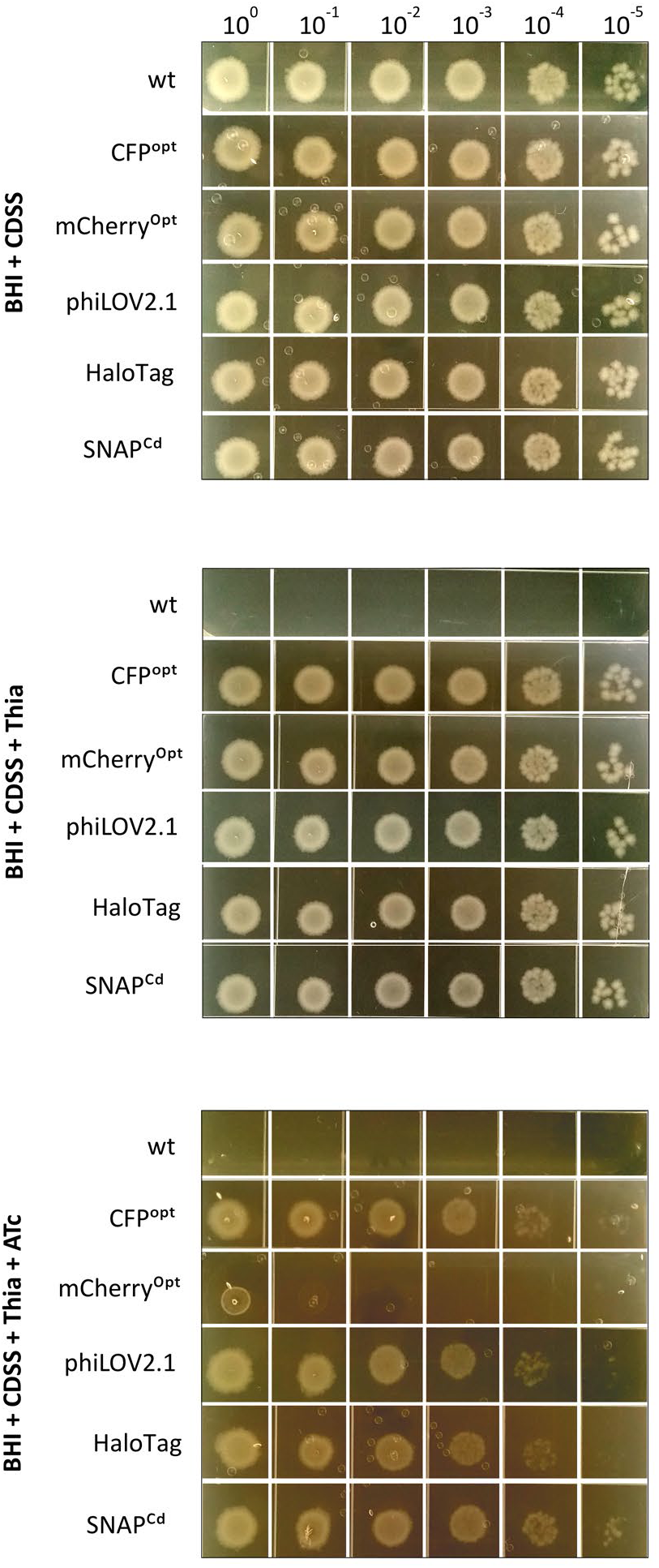
Viability is reduced upon expression of cytosolic mCherry^Opt^. Analysis of the viability of *C. difficile* 630Δ*erm,* and strains harboring different fluorescent systems: WKS1719 (CFP^opt^), AF312 (mCherry^Opt^), WKS1734 (phiLOV2.1), AF313 (HaloTag) and AF314 (SNAP^Cd^) by spotting aliquots of serially diluted cultures. Plates (BHI/YE + CDSS, BHI/YE + CDSS + thia and BHI/YE + CDSS + thia + 200 ng/mL ATc) were incubated for 24 hours. The image is representative of three independently performed assays.

### Signals from the cytosolic fluorescent systems are highly variable

To benchmark selected fluorescent systems as reporters for gene expression (i.e. expressed as cytosolic proteins), we compared characteristics of CFP^opt^, phiLOV2.1, HaloTag and SNAP^Cd^. We determined fluorescence intensity, percentage of expressing cells (at the single-cell level) as well as expression level in the population of cells through in-gel fluorescence (Fig. 6). To compare the performance of the fluorescent systems in *C. difficile,* the strains harbouring the different constructs where induced in exponential growth phase with 200 ng/mL ATc for 1 hour after which samples were collected for microscopy and in-gel detection. Fluorescence microscopy of the samples was performed at an approximate OD_600nm_ of 0.8, with appropriate filter settings to achieve a better signal to noise ratio and prevent bleed-through between channels (see Material and Methods). As for the intrinsic cell fluorescent analysis, no membrane dye was added, but nucleoid staining was performed to assess the influence on the chromosome. No differences on the nucleoid were observed for all fluorescent systems when compared to wt (Fig. 6).

**Fig. 2.**
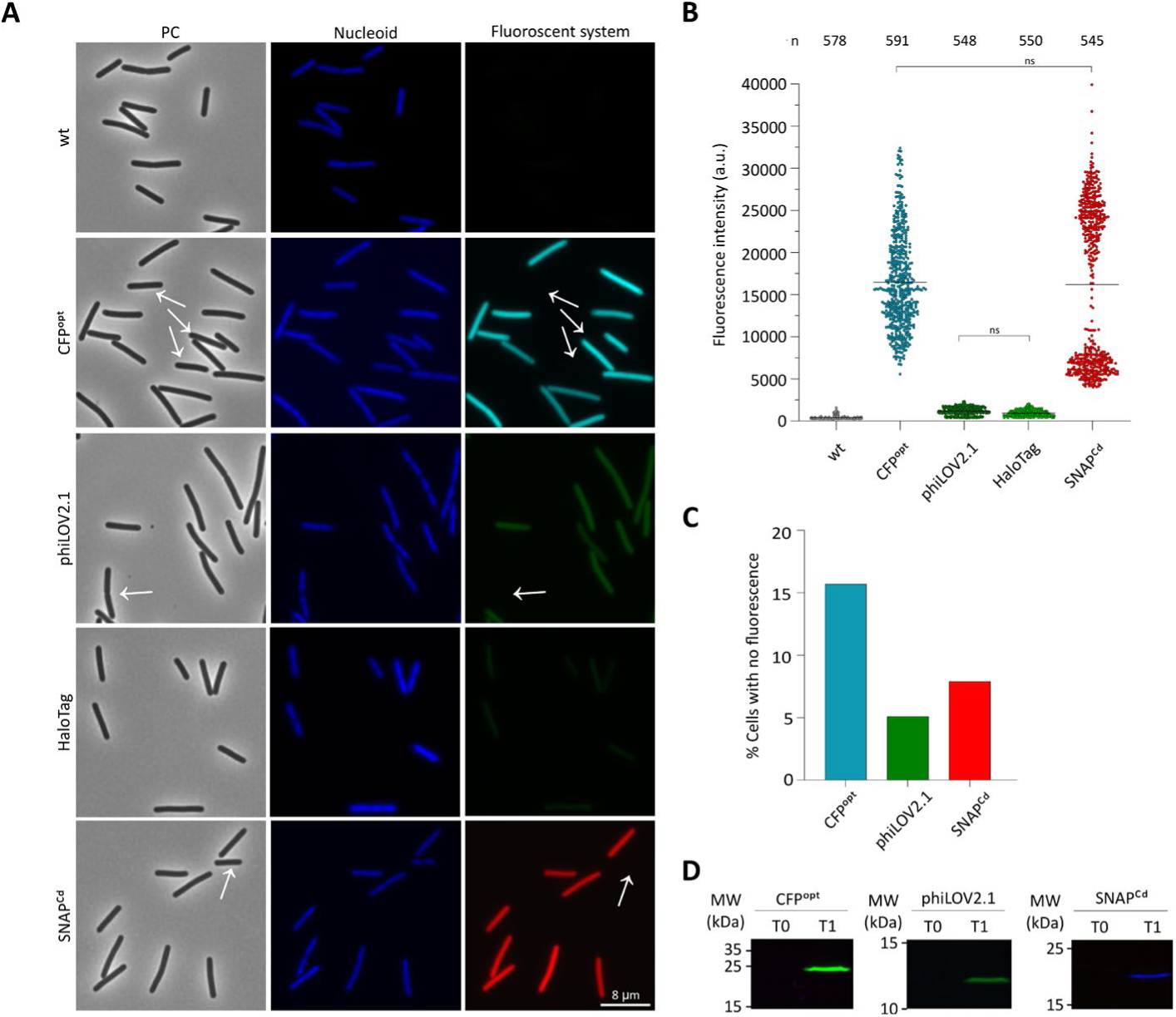
The cytosolic fluorescent systems of *C. difficile*. Analysis of *C. difficile* 630Δ*erm* and harbouring the vectors for the ATc-dependent overexpression of the cytosolic fluorophores, WKS1719 (CFP^opt^), AF312 (mCherry^Opt^), WKS1734 (phiLOV2.1), AF313 (HaloTag) and AF314 (SNAP^Cd^). **A)** Fluorescence microscopy of cells induced at mid-exponential growth phase in liquid medium with 200 ng/mL ATc for 1h. The effect was verified by three independent microscopy experiments. Cells were stained with DAPI dye for nucleoid visualization. Phase contrast (PC) and correspondent channel (see Material and Methods) are shown. Cells with no fluorescent signal are indicated with a white arrow. Because growth is asynchronous in these conditions, cells representing different cell cycles stages can be found. Scale bar = 8 µm. **B)** Mean longitudinal fluorescence intensity (a.u.) in the correspondent channel of the analysed *C. difficile* cells. Standard deviation is represented. Quantifications were performed using MicrobeJ from at least two biological replicates for each condition, the number of analysed cells (n) is indicated above the graph. *p<0.0001 by one-way ANOVA. **C)** Percentage of cells that do not contain detectable fluorescence in the different *C. difficile* strains. **D)** In-gel fluorescent analysis of the *C. difficile* strains samples before induction (T0) and 1 hour after induction (T1). Samples were run on 15% SDS-PAGE.

The CFP^opt^ protein has previously been codon-optimized for expression in low GC gram-positive bacteria [24, 38]. It should be noted that visualization of this fluorophore requires exposure of the sample to environmental oxygen, for fluorophore maturation. Without such exposure, overexpressing the CFP^opt^ does not lead to a detectable fluorescent signal (data not shown). Therefore, samples were exposed for 30 min to ambient oxygen outside the anaerobic cabinet. A bright fluorescent signal (16463 ± 5599 A.U., Fig. 6A-B) was observed in the majority of *C. difficile* cells, although ∼16% of cells had no visible fluorescent signal (Fig. 6A, white arrows, Fig. 6C). The high variability observed between the cells could also be due to the differences in oxygen exposure and accessibility in our setup. The use of fixed samples with overnight oxygen exposure as previously reported might overcome some of the variability we observe [24]. CFP^opt^ is also suitable for in-gel detection, as upon induction of CFP^opt^ expression, a signal is observed with an approximate molecular weight of 27 kDa, as predicted (Fig. 6D).

Similar to CFP^opt^, mCherry^Opt^ requires exposure to ambient oxygen. However, consistent with our previous observation of reduced viability upon induction of cytosolic mCherry^Opt^, induction of cytosolic mCherry^Opt^ in liquid culture led to cell death that prevented us from determining expression by fluorescent microscopy (data not shown).

In contrast to CFP^opt^ and mCherry^Opt^, phiLOV2.1 is a reportedly oxygen-independent fluorophore and has been previously proposed as a promising candidate for fluorescence in the green channel in *C. difficile* [28]. phiLOV2.1 is a flavin mononucleotide-based fluorescent protein that has been improved for enhanced photostability [27]. Under our conditions, however, overexpression of phiLOV2.1 resulted in very low intensity signals (1183. ± 462 A.U., Fig. 6A and B) but only 5% of cells did not exhibit any detectable fluorescence (Fig. 6A, white arrows, Fig. 6C). Overexpression of phiLOV2.1 was detected after 1-hour induction at the approximate molecular weight of 12 kDa (Fig. 6D). Previous reports on phiLOV2.1 expression used antifading compounds [28] that were omitted in this study, for consistency between fluorescent systems. The use of antifading could improve the usability of this fluorophore.

Another oxygen-independent tag is the HaloTag, which requires the addition of a ligand for visualization. Out of multiple ligands available, the Oregon Green substrate (green fluorescence) was used with 30 min incubation, similar to previous studies where the tag was fused to the HupA protein [33] (see also below). Very low levels of fluorescence were detected (943.9 ± 352.6 A.U., Fig. 6A-B). These results suggest very low intensity signals or no fluorescence signal from the HaloTag overexpression, which poses a problem when evaluating this fluorophore in the same channel that detects the intrinsic fluorescence of *C. difficile*. In addition, no clear overexpression was detected for the HaloTag construct 1 hour after induction (Fig. 6D), and our conditions, therefore, did not allow a proper assessment of this tag in *C. difficile*.

Similar to the HaloTag, the SNAP^Cd^ requires the addition of a ligand. Here, we employed the TMR substrate (red fluorescence) with incubation for 30 min, as this is most commonly used in published studies [31, 52, 53]. Visualization of SNAP^Cd^ overexpression after 1-hour induction shows a strong fluorescence signal in the red channel, but also a large variability in signal intensity (16178 ± 9394 A.U., Fig. 6A-B). The variability of the fluorescence signal observed may reflect either heterogeneous uptake of the substrate, or variable expression of the fluorophore. Only 8% of cells did not contain fluorescence (Fig. 6A, white arrows, Fig. 6C). Overexpression of SNAP^Cd^ in the culture was confirmed by in-gel fluorescence, with the presence of a band with the predicted molecular weight of 19.7 kDa (Fig. 6D).

Taken together our results suggest that not all fluorescent systems are equally suitable to evaluate gene expression at the single-cell level in *C. difficile*. CFP^opt^ and SNAP^Cd^ exhibit the highest fluorescent signal intensities and show no apparent toxicity when expressed in the cytosol.

### Fusion to fluorescent systems does not prevent the lethal effect of HupA overproduction

Next, we aimed to compare the different fluorescent systems as markers for protein expression and localization. We constructed modular vectors expressing the selected fluorescent systems (CFP^opt^, mCherry^Opt^, phiLOV2.1, HaloTag and SNAP^Cd^), fused to the C-terminus of the HupA protein [33] (Fig. 4). Thus, all constructs once again contained the same inducible promoter (P*_tet_*), identical ribosome binding sites (derived from the *cpw2* gene) [37], and the same GS-linker (Fig. 4, see Materials and Methods).

HupA protein is an essential bacterial chromatin protein in *C. difficile*. It has previously been shown that overproduction of HupA in *C. difficile* leads to lethal compaction of the chromosome [33]. This feature makes HupA an interesting candidate to study the effect of the different fluorescent systems, as any tag that would interfere with the functionality of the HupA protein, would likely affect the HupA-dependent decrease of cell viability.

To evaluate the effect of overexpression of the HupA fusion to the different fluorescent systems on the viability of *C. difficile* cells, we first performed a spot assay. The different *C. difficile* strains were spotted in 10-fold serial dilutions on BHI/YE agar with *C. difficile* selective supplement (CDSS), supplemented with thiamphenicol and anhydrotetracycline (ATc) when appropriate. All the strains grew indistinguishably until a dilution of 10^−5^ on medium lacking thiamphenicol and ATc (Fig. 7, top panel). In the absence of inducer but the presence of thiamphenicol, all strains also show clear growth up to a dilution of 10^−5^ (Fig. 7, middle panel) indicating that selection for the expression plasmid does not lead to reduced viability.

**Fig. 3.**
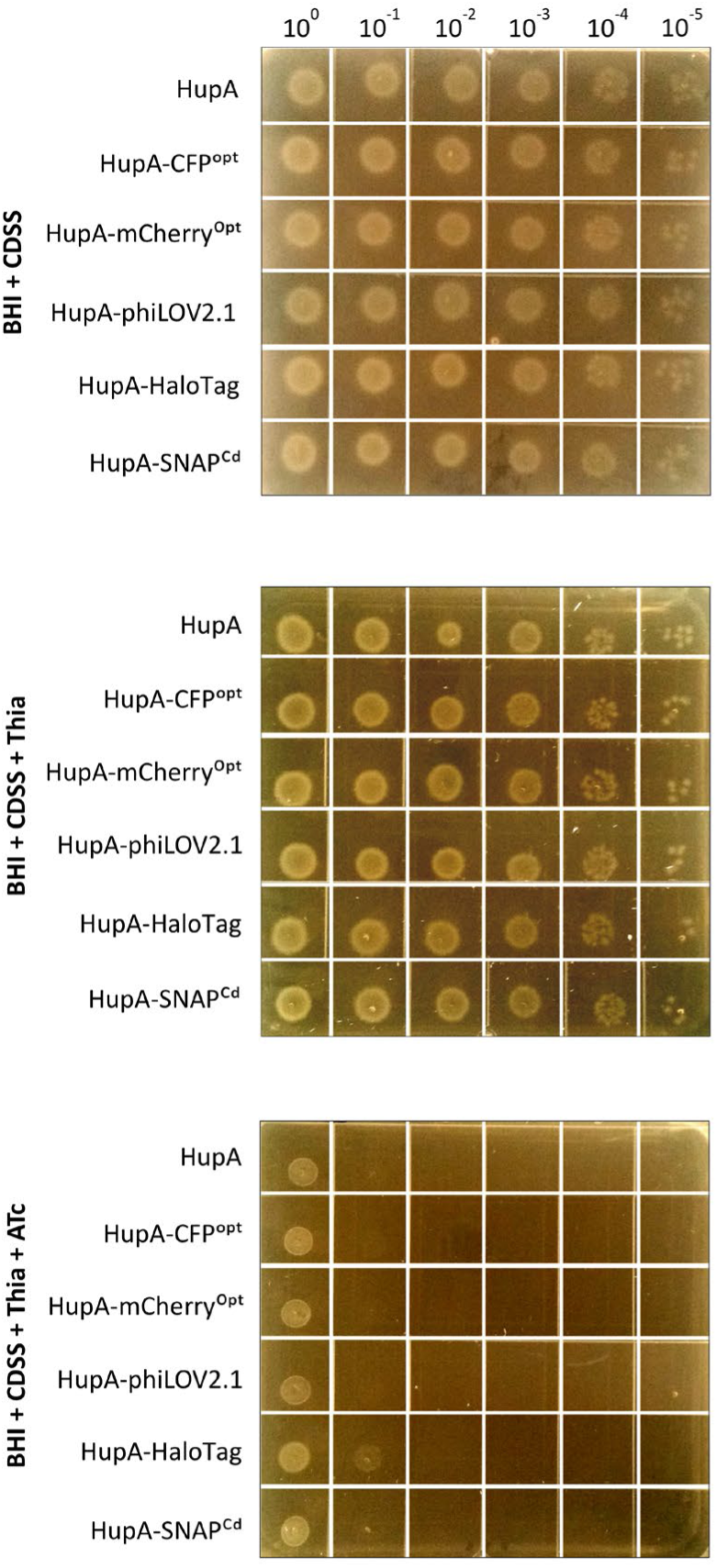
Tagging with fluorescent systems does not affect the reduction in viability associated with HupA overexpression. Analysis of the viability of *C. difficile* 630Δ*erm* harboring the different fluorescent systems: RD14 (HupA-CFP^opt^), RD15 (HupA-mCherry^Opt^), RD13 (HupA-phiLOV2.1), RD16 (HupA-Halotag) and RD17 (HupA-SNAP^Cd^) by spotting aliquots of serially diluted cultures. Plates (BHI/YE + CDSS, BHI/YE + CDSS + thia and BHI/YE + CDSS + thia + 200 ng/mL ATc) were incubated for 24 hours. The image is representative of three independently performed assays.

In line with previous results [33], we observed an approximate 4-log reduction in viability in the presence of ATc upon overexpression of non-tagged HupA. Notably, fusions to the C-terminus of HupA to the N-terminus of the fluorescent systems showed a similar reduction in viability (Fig. 7, lower panel). We repeatedly observed that a fusion of HupA to the HaloTag showed some limited growth at 10^−1^ (Fig. 7 and [33]), suggesting that the HaloTag exerts a modest effect on HupA functionality. As noted above, the HaloTag is the only fluorophore in our experiments that has not been codon-optimized for expression in *C. difficile*, and the defect may be the result of the atypical codon usage. In contrast to the cytosolic HaloTag, fusion to the C-terminus of HupA might minimize the impact of the lack of codon optimization.

HupA-mCherry^Opt^ overexpression shows no growth at 10^−1^ (Fig. 7, lower panel), similar to previously observed when expressing the cytosolic mCherry^Opt^ (Fig. 5, lower panel). However, in contrast to the cytosolic version, HupA-mCherry^Opt^ expression is conditioned by the HupA expression level, which by itself reduces cell viability. Thus, the reduction in viability observed could be a result of the mCherry^Opt^ toxicity or the HupA overexpression effect.

These results suggest that, with the possible exception of the HaloTag and/or mCherry^Opt^, the fluorescent systems do not affect HupA protein function.

### Fusions of the fluorescent systems to HupA localize to the nucleoid

Finally, we evaluated the performance of the fluorescent systems as tools to localize proteins subcellularly in *C. difficile*. If DNA binding of HupA is not affected, we expect all fusion proteins to localize to the nucleoid [33]. We also assessed the impact of the fusions on the HupA-mediated nucleoid condensation, which is a more subtle indication of HupA functionality, through staining of the nucleoid (Fig. 8A-B) [33]. In addition to HupA localization and function, we also quantified fluorescent intensity, percentage of expressing cells, and aggregate expression levels through in-gel fluorescence (Fig. 8B-E). To this end, strains harbouring the different fusion constructs were induced in exponential phase with 200 ng/mL ATc for 1 hour after which samples were collected for microscopy and in-gel detection.

**Fig. 4.**
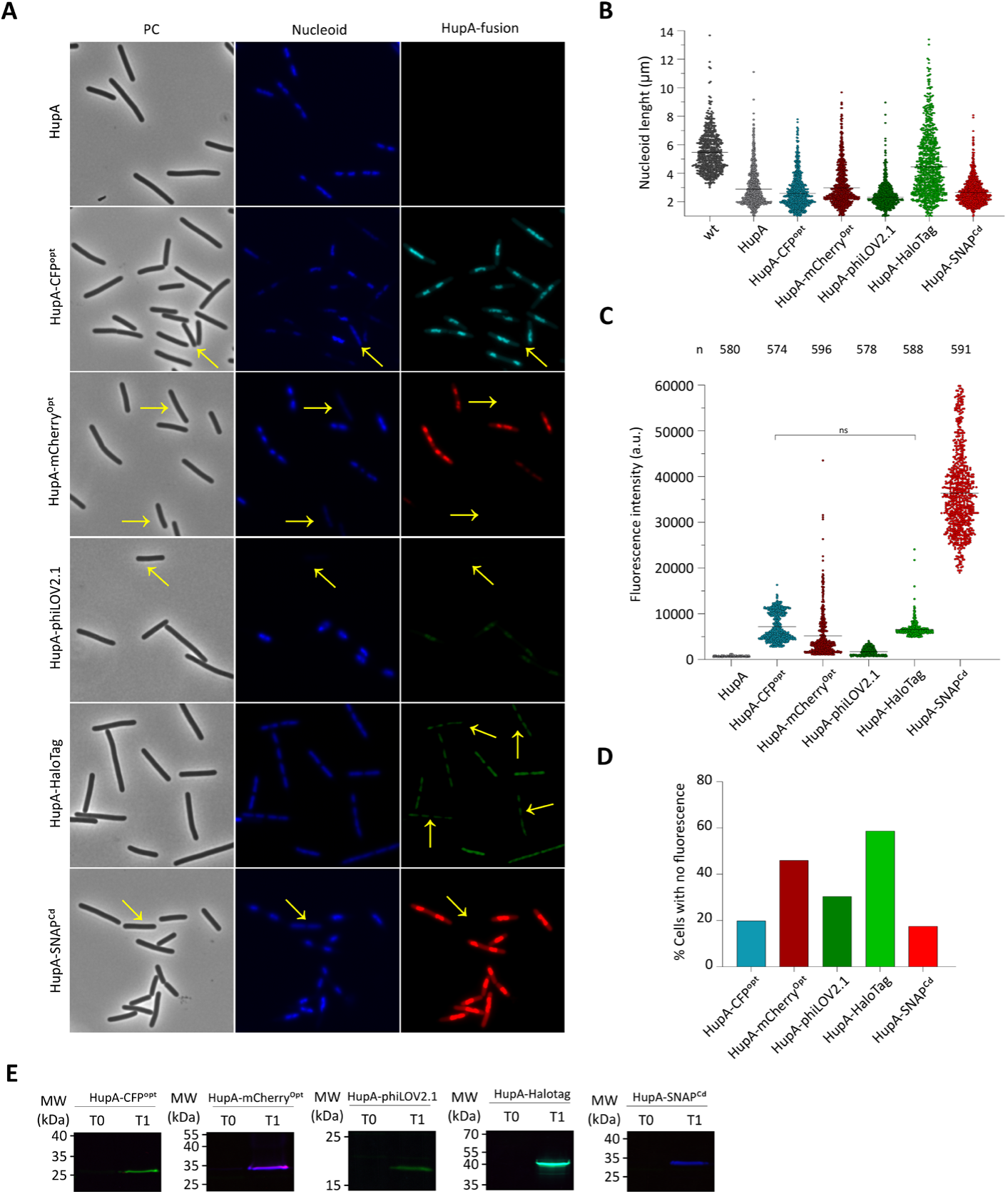
HupA tagged with different fluorescent systems localizes to the nucleoid. **A)** Fluorescence microscopy analysis of *C. difficile* 630Δ*erm* harbouring the vectors for the ATc-dependent overexpression of the cytosolic fluorescent systems, *C. difficile* strains AP106 (HupA), RD14 (HupA-CFP^opt^), RD15 (HupA-mCherry^Opt^), RD13 (HupA-phiLOV2.1), RD16 (HupA-HaloTag) and RD17 (HupA-SNAP^Cd^). Cells were induced at mid-exponential growth in liquid medium with 200 ng/mL ATc for 1h. The effect was verified by three independent microscopy experiments. Cells were stained with DAPI dye for nucleoid visualization. Phase contrast (PC) and correspondent channel (see Material and Methods) are shown. Cells with no expression are indicated with a yellow arrow. Because growth is asynchronous in these conditions, cells representing different cell cycles stages can be found. Scale bar = 8 µm. **B)** Mean nucleoid size (µm) of the analysed *C. difficile* cells. Standard deviation is represented. Quantifications were performed using MicrobeJ from at least two biological replicates for each condition, 500-600 cells were analysed per condition. *p<0,0001 by one-way ANOVA. **C)** Mean longitudinal fluorescence intensity (a.u.) in the correspondent channel of the analysed *C. difficile* cells. Standard deviation is represented and quantification was performed using MicrobeJ. **D)** Percentage of cells that do not contain detectable fluorescent signal in the different *C. difficile* strains. **E)** In-gel fluorescent analysis of the *C. difficile* strains samples before induction (T0) and 1 hour after induction (T1). Samples were run on 12% SDS-PAGE.

Overexpression of non-tagged HupA leads to a clear reduction of the nucleoid size (2.9 ± 1.3 µm, Fig. 8A-B) when compared to wt (5.5 ± 1.4 µm, Fig. 6A) consistent with previous observations [33]. Although some variation of DAPI staining is observed between the cells, it allowed for a straightforward analysis of the nucleoid sizes (Fig. 8). The fluorescent signal of different HupA fusions co-localize with the DAPI stained nucleoid, where compaction of the *C. difficile* nucleoid is observed in all the HupA fusions (Fig. 8A-B). Similar to the non-tagged HupA, the HupA fusions HupA-CFP^opt^ (2.6 ± 1.3 µm), HupA-mCherry^Opt^ (3.0 ± 1.5 µm), HupA-phiLOV2.1 (2.3 ± 1.0 µm) and HupA-SNAP^Cd^ (2.7 ± 0.9 µm) have a reduced nucleoid size, but the HupA-HaloTag exhibited a less pronounced effect (but still smaller nucleoid than wt) (4.5 ± 2.3 µm, Fig. 8A-B). As suggested above (Fig. 7) and in a previous study [54], there appears to be an interference of the HaloTag with HupA function leading to an overall lower reduction on the nucleoid size.

Overexpression of HupA-CFP^opt^ leads to lower fluorescence intensity values and lower variability between the cells (7183 ± 2998 A.U., Fig. 8A and C) when compared to the cytosolic version (16463 ± 5599 A.U., Fig. 6A-B). Nevertheless, the number of cells with no fluorescent signal for the HupA-CFP^opt^ fusion was similar to the number of cells without a detectable signal for the cytosolic CFP^opt^ (20% vs 16%, Fig. 8D and 6C). The cells with no fluorescent signal did not exhibit condensation of the chromosome, consistent with no HupA-CFP^opt^ expression in these cells (Fig. 8A, yellow arrows). In-gel fluorescence analysis shows a band with the approximate molecular weight of 37 kDa, as predicted (Fig. 8E), although lower levels of expression of the HupA translational fusion are observed when compared with the free cytosolic CFP^opt^ (Fig. 6D).

The expression of HupA-mCherry^Opt^ did not appear to be toxic for *C. difficile* cells, as we were able to grow and image *C. difficile* cells expressing HupA-mCherry^Opt^. In a similar manner to HupA-HaloTag, expression of mCherry^Opt^ fused to HupA does not show the same toxic effect as the cytosolic counterpart, possibly due to the effect of the HupA fusion on the translational efficiency in *C. difficile*. Expression of HupA-mCherry^Opt^ exhibit a clear fluorescent signal co-localizing with the nucleoid, however low levels of fluorescence were observed (5174 ± 4957 A.U, Fig. 8A and C). Moreover, a high cell-to-cell variability is present for HupA-mCherry^Opt^ and 46% of the cells did not exhibit quantifiable fluorescence (Fig. 8A, yellow arrows, Fig. 8D). However, a clear signal of expression on the whole population is observed with the presence of a band with the predicted molecular weight of approximately 37 kDa (Fig. 8E). Interestingly, we initially imaged the HupA-mCherry^Opt^ fluorophore also with a different filter set (mCherry, Leica n. 11532447, ex: 580/20, dichroic mirror: 595, em:630/60), which would be theoretically more suitable for this fluorophore, but very low intensity values were obtained when compared with the filter ultimately used (Y3, Leica n. 11504169, ex: 545/25, dichroic mirror: 565, em:605/70; data not shown). Both of HupA-CFP^opt^ and HupA-mCherry^Opt^ fluorescent signals could potentially be enhanced by longer oxygen exposure and oxygen accessibility [24, 25].

Unexpectedly, no clear fluorescence signal was observed for HupA-phiLOV2.1 overexpression when the cells imaging was performed immediately after sampling (Fig. 9). However, a clear reduction in nucleoid size was observed suggesting overexpression of HupA-phiLOV2.1 (Fig. 9). Indeed, expression of the protein was confirmed by the presence of the band with predicted molecular weight, at the moment of sampling (∼24 kDa, Fig. 8E). We noted, however, that anaerobic incubation of our samples for 30 min, similar to the substrate incubation required for the SNAP^Cd^ and HaloTag, led to a detectable HupA-phiLOV2.1 fluorescence signal (6600 ± 1529 A.U., Fig. 8A and C). However, 30% of the cells did not express HupA-phiLOV2.1 as no fluorescence and HupA-induced chromosome condensation was observed (Fig. 8A, yellow arrows, Fig. 8D).

**Fig. 5.**
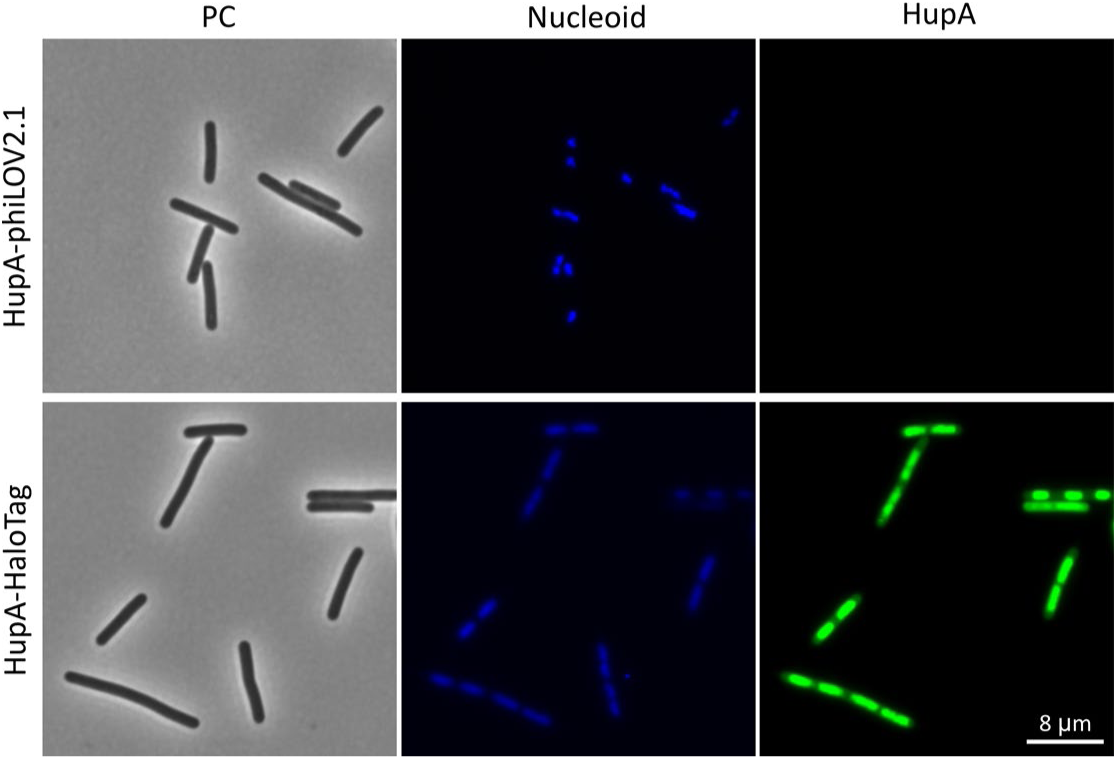
HaloTag visualization with antifading and immediate imaging of phiLOV. Fluorescence microscopy analysis of *C. difficile* 630Δ*erm* and harboring the vectors for the ATc-dependent overexpression of the cytosolic fluorescent systems, *C. difficile* strains RD13 (HupA-phiLOV2.1) and RD16 (HupA-Halotag). Cells were induced at mid-exponential growth in liquid medium with 200 ng/mL ATc for 1h. Cells where stained with DAPI dye for nucleoid visualization. Immediate imaging of HupA-phiLOV2.1 after DAPI staining. Use of antifading reagent for HupA-HaloTag microscopy visualization. The effect was verified by three independent microscopy experiments. Phase contrast (PC) and correspondent channel (see Material and Methods) are shown. Because growth is asynchronous in these conditions, cells representing different cell cycles stages can be found. Scale bar = 8 µm.

HupA-HaloTag overexpression has been previously described in *C. difficile* and the results obtained here are consistent with that study [33]. Overexpression of HupA-HaloTag results in a clear fluorescent signal in the green channel (6600 ± 1529 A.U., Fig. 8A and C). However, a very high percentage of cells did not present any detectable fluorescence or lower than the defined threshold settings (59%, Fig. 8A, yellow arrows, Fig. 8D, see Material and Methods). Notably, lower levels of fluorescence were observed in the current study when compared to previous data [33]. To confirm the influence of differences in the experimental set-up in this study we performed additional experiments with the addition of anti-fading reagent as previously reported (Fig. 9) [33]. Indeed, the addition of anti-fading agent to the microscopic preparation resulted in a 1-log increase in the fluorescent signal (27952 ± 12955 A.U.) and better definition of the signal-to-noise ratio, resulting in better visualization of the fusion protein (Fig. 9). It should be noted, however, that different substrates are commercially available for use with the HaloTag and the use of substrates with different spectra could potentially affect signal intensities too. In contrast with free cytosolic HaloTag (Fig. 6), induction led to the presence of the ∼45 kDa band, consistent with the theoretical MW of HupA-HaloTag (Fig. 8E). Poor translation due to a non-optimized codon-usage as observed for the free HaloTag, is ameliorated by the fusion to the HupA protein.

Induction of HupA-SNAP^Cd^ overexpression results in a clear fluorescence signal in the red channel with the highest intensity values (36379 ± 8882 A.U., Fig. 8A and C) when using the TMR substrate. Low cell-to-cell variability between cells was observed for the HupA fusion when compared to the free cytosolic version of SNAP^Cd^ (Fig. 6B). Compared to the other fluorescent systems only 18% of cells exhibit no fluorescence (Fig. 8A, yellow arrows, Fig. 8D), outperforming the other fluorescent systems for HupA visualization. As expected, overexpression led to the presence of a band ∼30 kDa, consistent with the predicted MW of HupA-SNAP^Cd^ (Fig. 8E).

Taken together, these results suggest that all the different fluorescent systems can be successfully employed in *C. difficile* to evaluate protein expression and localization. Yet, there were major differences between specific characteristics of the fluorescent systems, that may limit the usefulness for certain applications.

## Discussion and Conclusion

### The source of autofluorescence in *C. difficile*

Autofluorescence, mainly in the green spectrum, in *C. difficile* has been a distinct disadvantage for fluorescence microscopy, leading to a search of fluorescent systems that overcome the interference of the intrinsic fluorescence, for use in gene expression and protein localization studies. In this study, we sought to better understand *C. difficile* autofluorescence and the differences observed in autofluorescence between different studies, with the ultimate goal to sidestep this problem in live-cell microscopy.

Microscopy of *C. difficile 630Δerm* cells during different stages of cell growth allowed us to identify that the autofluorescence is influenced strongly by the growth stage of the sampled culture (Fig. 1). At the onset of the stationary growth phase the fluorescence signal significantly increased. *C. difficile* growth is non-synchronous and at this stage different subpopulations of cells are present, likely induced by nutrient depletion and the onset of sporulation [55]. Indeed, several studies where fluorescence microscopy is used for the study of *C. difficile* developmental processes show a highly variable autofluorescence depending on the cell growth stage, where low levels of green fluorescence are detected in exponential phase and higher levels in stationary [31, 33].

Growth phase-dependent autofluorescence has also been observed for organisms such as *E. coli* or *B. pumilus* [30, 41]. In *E. coli*, ribo(flavin) compounds have been identified as the main components responsible for the green fluorescence [30]. A link between autofluorescence and oxidative stress has been proposed, as the pigment responsible for the autofluorescence was found to be influenced by environmental cues, such as H_2_O_2_ [41].

Flavoproteins act as electron transfer proteins and mediate redox reactions, for instance in the metabolism of carbohydrates, oxidative stress response or even in the extracellular environment (Garcia-Angulo 2017). In *Streptococcus pneumoniae*, the flavin reductase FlaR contributes to the resistance to H_2_O_2_ [43]. The expression of genes involved in the synthesis and uptake of flavins is regulated through FMN-dependent riboswitches [56, 57]. Flavodiiron proteins in anaerobic bacteria such as *C. difficile* are involved in oxidative stress, allowing the cells to cope with the presence of oxygen or reactive oxygen species [44].

In *C. difficile,* an association of flavodiiron proteins with the regulators of the general stress response [45, 46] and sporulation [58] has been observed. However, when evaluating the autofluorescence of strains lacking the *sigB* gene (Δ*sigB*) or carrying a disrupted *spo0A* gene (*spo0A*::CT) no significant difference from wild type cells was observed (Fig. 2 and S1). Nevertheless, exposure to environmental oxygen results in a significant increase in the autofluorescence in *C. difficile* (Fig. 3), suggesting a mechanism independent from the general stress response or sporulation pathways.

In the course of our experiments, we observed that *C. difficile* expressing the non-tagged HupA protein demonstrated higher levels of fluorescence in the green channel than expected for the growth stage. We note, however, that these samples were imaged after all other samples towards the end of the microscopy sessions. Thus, they were subjected to the waiting times and possibly some environmental oxygen which may have entered the agar patches, despite our efforts to prevent this, which might explain the increased fluorescence.

In a previous study, exponentially growing *C. difficile* cells expressing mCherry^Opt^ were subjected to prolonged exposure to environmental oxygen, required for fluorophore maturation, and exhibited high levels of autofluorescence [24, 25], consistent with our observations. Additionally, HupA-induced chromosome condensation and/or the burden of overproduction could also affect or potentiate the autofluorescence of *C. difficile* cells.

Taken together with the previous observations, our data suggest that *C. difficile* autofluorescence might result from direct oxidation of specific cell components, which may vary in abundance dependent on growth phase or cell cycle stage. The identification and presence of (ribo)flavins and flavoproteins is largely unexplored in *C. difficile* and might be the reason for oxidation-dependent autofluorescence.

As the pathways governing autofluorescence are unknown, such fluorescence could limit the study of specific pathways. Additionally, we note that studies of *C. difficile* in stationary growth phase or under certain stress conditions could demonstrate significant autofluorescence, potentially limiting the use of green-spectra fluorescent systems. It is noteworthy that autofluorescence as observed here showed high levels of cell-to-cell variability that could interfere with normalization of fluorescence intensities. Thus, careful use of green spectra fluorescent systems is recommended and further studies are required to identify the compounds responsible for the fluorescence observed and their possible role on the bet-hedging strategy of *C. difficile*.

### Possible interference of autofluorescence with fluorescent systems analysis

Fluorescent microscopy applications are very wide [59, 60]. The choice of imaging platform and filter sets is crucial, as this can have a large effect on the accuracy of the observation. For this study, the microscopy conditions and filter sets are listed in the Material and Methods section. Our results should be interpreted in light of the set-up used, and may not yield the same results when replicated by others using different methods.

There is a wide range of filter sets that allow for multiple fluorescent systems visualization, however fluorescence channel bleeding can be a problem when imaging multiple fluorescent systems or dyes on wide-field fluorescence microscopy. In our microscopy settings bleed-through between the cyan (CFP) and green channels (Y3-Alexa 488/L5) could be detected (data not shown). The CFP^opt^ fluorophore is imaged at an emission wavelength that can also capture the *C. difficile* autofluorescence (approximately 480-550 nm) as previously noted by others [25]. Nevertheless, with sufficient CFP^opt^ expression and maturation, its signal can be clearly distinguished over the background signal.

Both phiLOV2.1, and HaloTag with the Oregon Green substrate exhibit fluorescence in the green spectrum (emission wavelength 500-570 nm), similar to the wavelengths of the intrinsic cell fluorescence. As for CFP^opt^, visualization over the mean background fluorescent signal of phiLOV2.1 and Halotag was possible. However, it is not possible to correct the background at the single-cell level when fluorescent systems and autofluorescence occur in the same channel. Thresholding is possible by calculating average values for the autofluorescence background but does not allow for background correction at the single-cell level. Fluorescent systems with fluorescence in these spectra are therefore not suitable for studying low levels of expression.

Though in this study we used measurements of intrinsic fluorescence to threshold fluorescence levels, this can be challenging due to cell-to-cell variation in both background and fluorescent systems-related fluorescence. Thus, the use of fluorescent systems with spectra that (partially) overlap with autofluorescence, requires careful analysis with background correction performed ideally at the single-cell level, and could potentially be minimized by the use of filter sets with a narrower bandwidth.

Recently, another limitation when imaging green spectra fluorescent systems, which rely on the excitation in the violet or blue channels (400-470 nm), was identified. It has been shown in different organisms, like *B. subtilis,* that cell growth is arrested when excited in violet and blue channels [61], which significantly reduces the use of CFP^opt^ fluorophore or even DAPI on live-cell imaging. However, the influence of the excitation wavelengths on *C. difficile* cell metabolism is still unexplored and outside the scope of your study.

### *C. difficile* fluorescent systems

Despite the challenges presented by autofluorescence, it did not restrict our further studies on the fluorescent systems, as the microscopy experiments presented here were conducted on exponentially growing cells with very limited oxygen exposure, except when needed for maturation.

In this study, we aimed to compare the performance of different fluorescent systems used in *C. difficile*. Modular vectors, using uniform expression signals, were created for the different fluorescent systems which should eliminate differences resulting from differences in native expression signals (Fig. 4). Furthermore, only exponentially growing cultures were sampled in a similar manner for all fluorescent systems (see Material and Methods). These steps, however, do not completely eliminate cell-to-cell variation, as a *C. difficile* culture demonstrates non-synchronous growth and different cell growth stages. Additionally, it is further potentiated by the heterogeneous expression of the fluorescent systems, plasmid copy number or gene expression noise [62–64].

We observed distinct differences between the fluorescent systems CFP^opt^ [24], mCherry^Opt^ [25], phiLOV2.1 [28], HaloTag [33] and SNAP^Cd^ [31] in evaluating the possible use for gene expression (cytosolic, Fig. 5-6) as well for as for protein localization and functionality studies (HupA-fused, Fig. 7-9). The most notable difference relates to the requirement of oxygen or a fluorescent substrate for visualization and therefore some aspects will be discussed in this context below, and are summarized in Table 5.

**Table 5.**
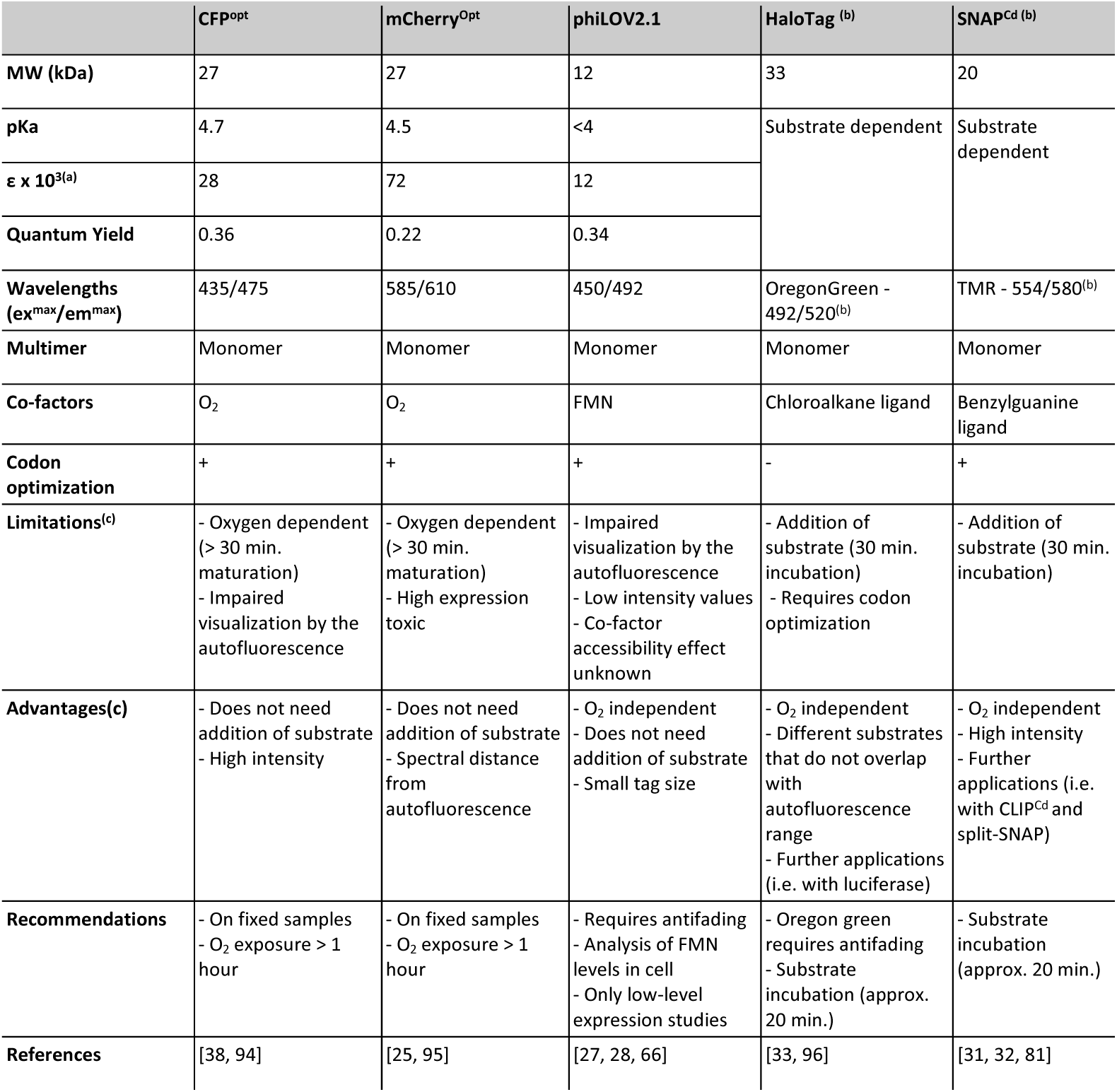
Overall properties and applicability in *C. difficile* of the different fluorescent systems used in this study. ^(a)^ Extinction coefficient (M^−1^ cm^−1^), ^(b)^ Different substrates are available, ^(c)^ Under the microscopy conditions used.

### The oxygen-dependent fluorophores CFP^opt^ and mCherry^Opt^

CFP^opt^ and mCherry^Opt^, require exposure to the environmental oxygen for fluorophore maturation, which is a major limitation, as oxygen exposure could lead to potential artefacts by affecting cell integrity and growth characteristics. Furthermore, the contribution of oxygen to autofluorescence, as observed in this study, could impair the proper analysis of gene expression and protein localization. Thus, these fluorophores are unsuitable for live-cell imaging and should only be performed on fixed cells.

We also observed that under our conditions expression of cytosolic mCherry^Opt^ was highly toxic to the cells. As the construct is codon-optimized it is unlikely that codon bias explains this phenomenon. Previous reports of lysis in non-fixed mCherry^Opt^ expressing cultures were attributed to 1-h oxygen exposure before visualization [25], but in our spot-assays, cells were not exposed to oxygen. Negative effects of mCherry as a reporter have been previously reported, where cells expressing mCherry were abnormally shaped compared to cells expressing GFP at the same locus of *Schizosaccharomyces pombe* [65]. However, when fused to HupA, expression of mCherry^Opt^ did not lead to evident cellular defects. Our results suggest that high-level expression of mCherry^Opt^ alone is toxic in *C. difficile* cells, limiting the applications of this fluorophore as a gene expression reporter.

### The oxygen-independent fluorophore phiLOV2.1

In theory, many of the drawbacks of the oxygen dependent fluorophores discussed above could be eliminated by the use of a FMN-based fluorescent protein, as FMN is believed to be available intracellularly and maturation of the fluorophore does not require molecular oxygen [66, 67]. Of this family of proteins (that also includes EcFbFP and iLOV, for instance) [27, 66, 68, 69], phiLOV2.1 has been used in *C. difficile* for the localization of cell division and flagellar proteins, FtsZ and FliC [28]. However, in your study, unexpectedly, we could only detect very low levels of phiLOV2.1 expression when expressed either by itself or fused to the HupA protein (Fig. 6 and 8). This may in part be attributed to the use of anti-fading compounds in previous work [28], whereas we, for consistency in our analyses, did not use such a compound when evaluating phiLOV2.1. Therefore, anti-fading compounds might improve the usability of this fluorophore, as we have also observed for HaloTag. However, the use of anti-fading compounds can impair live-cell microscopy, and new approaches might be required, as the addition of antifading agents in the culture medium [70].

Although phiLOV2.1 was engineered for higher photostability and does not require oxygen for maturation of the fluorophore, it comprises a LOV domain which requires the binding to the cellular FMN [27]. Though the reasons for low-level fluorescence of phiLOV2.1 require further investigation, it seems possible that intracellular FMN-levels are insufficient for high-level fluorescence and that FMN pools could vary depending on experimental conditions. In *E.coli*, it was shown that phiLOV2.1 is not suitable for protein localization studies of proteins present in the periplasmic space [71]. In *C. difficile* the use of phiLOV2.1 to localize extracellular protein domains has been shown, suggesting that FMN is also exported to the environment or co-translocated with the phiLOV2.1 protein [28]. High expression levels, such as in this study, can lead to a rate of protein synthesis that can potentially exceed the available intracellular FMN. The availability of intracellular FMN, as well as the levels of FMN required for cellular metabolism, have not yet been determined and could affect this fluorophore.

We observed a slow increase in fluorescence signal for phiLOV2.1 constructs over time and others have made similar observations [28]. These results can be explained by the FMN availability or an unknown process required for maturation of the fluorophore in *C. difficile*. Consistent with this notion, we saw clear compaction of the nucleoid, suggestive of HupA-phiLOV2.1 overexpression, prior to detecting the fluorescent signal (Fig. 8A and 9). phiLOV2.1 was codon-optimized for *C. difficile*, but maturation of phiLOV2.1 might still be sub-optimal, as codon bias can not only affect the rate of protein synthesis but also protein folding efficiency, for instance [72].

Overall, because of the possible drawback of limiting FMN pools and slow maturation, we feel that the use of phiLOV2.1 for live-cell imaging is suboptimal and future use of the phiLOV2.1 in *C. difficile* requires an in-depth study of FMN levels throughout growth and possible engineering for faster maturation in *C. difficile*.

### Substrate-dependent tags as fluorescent systems in *C. difficile*

The limitations of both the oxygen-dependent and phiLOV2.1 as discussed above indicate that substrate-dependent fluorescent systems may be the best option for *C. difficile* live-cell imaging. Indeed, the availability of a wide diversity in substrates with different wavelengths (including those that do not overlap the autofluorescence spectrum), presents a significant advantage. Nevertheless, we observed some limitations.

First, we did not observe expression of the cytosolic HaloTag in *C. difficile* cells, whereas expression could be detected when fused at the C-terminus of HupA, as previously reported [33]. The HaloTag construct was not codon-optimized for expression in *C. difficile*, in contrast to the other analysed fluorescent systems. The original HaloTag has a [G+C] content of approximately 60%, which is substantially higher than the average [G+C] content for *C. difficile* (29%) [35]. This could potentially impair transcription and translation in *C. difficile*, as seen for other fluorescent systems when expressed in low-[G+C] gram-positive microorganisms [38, 72].

SNAP^Cd^ has been extensively used for protein localization and gene expression studies in *C. difficile* [31, 47]. The data from this study support the use of SNAP^Cd^ as fluorescent systems in *C. difficile* as it exhibits high fluorescent signal intensities when expressed in the cytosol or fused to HupA. Due to the hight intensity of the fluorescent signals, it was possible to clearly observe the widespread fluorescent signals.

Our study did reveal cell-to-cell variation in fluorescent signals (Fig. 6 and 8). Though these may reflect actual differences between cells, it is important to take into account that these could also arise due to the variability in the reaction that links the substrate to the protein tag, or to the heterogeneous uptake (which may vary between substrates) [73]. Additionally, providing sufficient substrate in a well-mixed system imposes significant technical challenges.

### The drawbacks in *C. difficile* microscopy and future perspectives

At the moment, the use of substrate-dependent fluorescent systems seems most promising for live-cell imaging of *C. difficile*. SNAP^Cd^ and HaloTag exhibit high fluorescent signal intensities when fused to HupA and due to the wide variety in substrates are currently the most promising candidates for live-cell imaging. The major drawbacks of HaloTag can probably be overcome by codon-optimization.

Several studies have used SNAP^Cd^ and HaloTag in combination with other fluorescent systems [32, 74]. For instance, previous research has shown the use of dual labelling experiments with SNAP^Cd^ and CLIP^Cd^ in *C. difficile* [32]. CLIP is a derivative of the SNAPtag protein, with engineered specificity to different substrates [75]. The different substrate availability and the specificity for the tags allow independent and specific labelling of both fluorescent systems in the same cell. For example, SNAP^Cd^ and CLIP^Cd^ were successfully used to assess σK and σG-dependent expression in different cellular compartments during sporulation [32].

Substrate-dependent tags offer also other interesting possibilities, such as super-resolution microscopy and combination with luminescence [60, 76, 77]. HaloTag and SNAPtag have been successfully used in combination for dual-colour imaging, to unravel the spatial and temporal organization of SPI1-T3SS and SPI4-T1SS, components of the secretion system, in *S. enterica* with STORM. By single-molecule tracking, a fusion protein of the HaloTag with MukB, a protein involved in chromosome segregation within the Structural Maintenance of Chromosomes (SMC), has been shown to achieve higher speed than a fusion protein with the mCherry, producing more reliable single-molecule localization data [78]. Finally, the HaloTag can be used in combination with the NanoLuc luciferase in a so-called NanoBRET assay, as reported for eukaryotic cells [79]. Nanoluc (Luc^opt^ for *C. difficile* use) is already extensively used in *C. difficile* [33, 54] and the combination with the HaloTag could provide a valuable tool for further applications, such as analysis of *in vivo* dynamics of protein interactions. Protein-protein interactions also be investigated using a split-SNAP or split-CLIP approach [80]. The SNAPtag is split into two domains, that can interact in close proximity to form a functional protein capable of binding the substrate and emit fluorescence [80]. In *C. difficile*, the system has been used to assess the interaction of the SpoIIQ and SpoIIAH complex [81].

Of note, there are several fluorophores (https://www.fpbase.org/), some of which have been used in *Clostridium* species, that have not been explored in this study, but that nevertheless might be useful reporters for *C. difficile* studies. Other oxygen-dependent fluorophores such as tdTomato, have been used in *C. difficile* as well [82]. The substrate-dependent fluorophore FAST has been used as a reporter protein for several genes involved in metabolism and allowed subcellular localization of the cell-division protein ZapA in *C. acetobutylicum* [83]. Recently, it was engineered (splitFAST) to assess the dynamics of protein-protein interactions in vivo, displaying a successful complementation assay [84]. UnaG, that has been reported to be highly fluorescent under anaerobic conditions [26, 29], has been unexplored in *C. difficile* to date. Despite the possibilities, these proteins may come with their limitations. For instance, proteins such as tdTomato are expected to suffer from the same limitations as CFP^opt^ and mCherry^opt^, (split)FAST proteins still require the addition of a substrate, and it has been reported that UnaG is poorly expressed unless fused to a tag to increase solubility [26]. Thus, the search for a co-factor and oxygen-independent fluorescent systems is still of interest to those studying anaerobic microbiology.

In our study, all fusions of the fluorescent systems to the C-terminus of the HupA protein successfully localized to the nucleoid. However, our study has two major limitations: the use of a plasmid-based overexpression system and assessing the functionality of the protein by nucleoid size reduction only. Overexpression in *C. difficile* may impose a metabolic burden, potentially increasing the intrinsic fluorescence, reducing the usability of the green-spectrum fluorescent systems. It may also result in localization patterns that are different from those observed with native levels of protein [85]. Moreover, overexpression in *C. difficile* is heterogeneous and the HupA-mediated reduction of the nucleoid size could potentially also impair the accessibility of the fluorescent systems’s co-factors, further reducing the signal intensities and increasing cell-to-cell variability. To address the overexpression limitation, the fusions could be introduced at the native genomic locus, for example using an Allele Coupled Exchange (ACE) [86] or CRISPR-Cas system [87].

It is also important to consider that the use of tags on the protein of interest can affect the function and/or localization of the protein, as also suggested by the slightly affected HupA function of the HupA-HaloTag construct (Fig. 8). We did not assess N-terminal fusions of HupA to the fluorescent systems. However, a careful comparison of phenotypes of tagged and non-tagged proteins at both the C- and N-terminus should be analyzed to rule out impaired functionality [17]. For these and other proteins, assessment of the effect of tag on the protein function and stability is an important step for achieving optimal microscopic conditions.

Several other factors contribute to the wide applicability of fluorescence microscopy, such as the labelling by fluorescent molecules or dyes, and the innovation of the technical platforms for imaging anaerobes. For maximal success in subcellular localization studies, tagged proteins should be combined with fluorescent stains of nucleic acids and lipids. To this end the nucleoid stain DAPI, also in this study, or the membrane dye FM4-64 have been widely used [31, 88]. Some of these compounds are summarized in Table 6. The advantages and limitations of each of these stains is outside of the scope of the present study, but it is important to note that the effects of these dyes are not yet fully explored and understood for *C. difficile*.

**Table 6.** Membrane and nucleoid dyes used in *C. difficile*.

|  | Staining | References |
| --- | --- | --- |
| MitoTracker Green FM probe | Membrane | [32, 97] |
| FM4-64 dye | Membrane | [31, 97] |
| DAPI stain | Nucleoid | [31, 33] |
| Hoechst stain | Nucleoid | [98] |
| HADA fluorescent D-amino acid | Membrane | [52] |

Besides addressing the limitations of fluorescent systems, the development of live-cell fluorescent microscopy for anaerobic bacteria also requires technical innovations. To fully explore *C. difficile* physiology through live-cell imaging, an anaerobic environment suitable for single cell analysis, without the need of sampling, is necessary. Several incubation systems are available for the different commercially available microscopes, that would allow recreating the environmental conditions for *C. difficile* growth, but ideally, the microscope should be placed inside an anaerobic cabinet. Recently, Courson *et al*. used so-called rose chambers for live-cell microscopy of *C. difficile* cells to measure the cell motility [89]. However, we observed that rich culture mediums often have compounds that interfere with the fluorescence microscopy (data not shown). Further experiments are required to define optimal media for the continuous live visualization of fluorescence in *C. difficile* cells [89]. Additionally, when using substrate-dependent fluorescent systems (SNAP^Cd^ and/or HaloTag) or other co-factors (such as FMN), these compounds need to be (continuously) fed into the live-cell imaging system, to ensure fluorescence of newly synthesized protein. The development of an anaerobic microfluidics system with regulable inlet valves would thus be advantageous in *C. difficile* microscopy [13, 90–92]. Developments of technical aspects of anaerobic fluorescence microscopy aid in overcoming some of the fluorescent systems limitations and would substantially enhance the usability of fluorescence microscopy for *C. difficile* studies.

In sum, the research here highlights key characteristics of existing fluorescent systems and identifies key areas for development in order for *C. difficile* to step into the spotlight of live-cell fluorescent microscopy.

## Acknowledgements

Work is supported by a Vidi Fellowship to W.K.S. (864.10.003) from the Netherlands Organization for Scientific Research (NWO) and a Gisela Thier Fellowship to W.K.S. from the Leiden University Medical Center. We thank Dr. Gillian Douce, Glasgow University, for kindly providing the phiLOV2.1 plasmid; Dr. Craig Ellermeier for kindly providing the mCherry^Opt^ containing vector pDSW1728; Dr. Adriano Henriques, ITQB-UNL, for kingly providing the pFT46 for the SNAP^Cd^ expression; and Dr. Jeroen Corver for helpful discussions.

## Author contributions

A.F., R.D., B.W. and W.K.S contributed to construction of the vectors and strains used in this study. A.P. conducted the sample preparation, microscopy and data analysis in this study. A.P. and A.F. conducted the expression analysis through in-gel detection. A.P. and W.K.S designed the study and wrote the manuscript with the contributions of all authors.

## Supplemental Figures

**Fig. S1.**
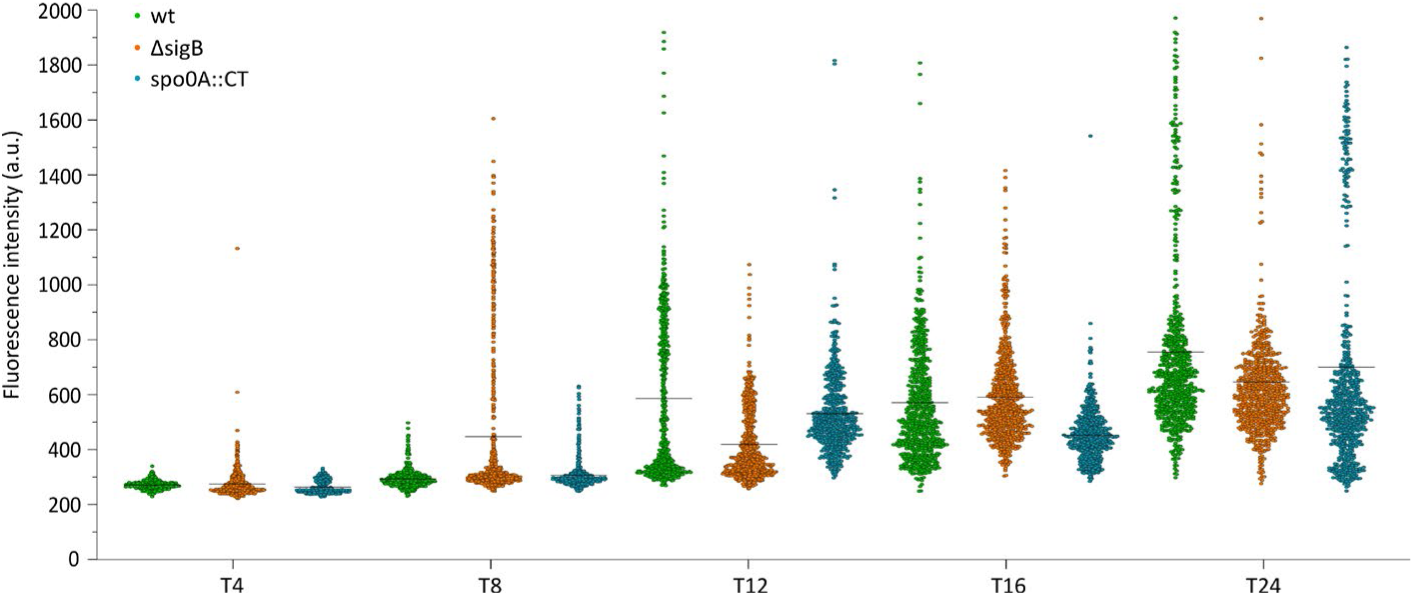
Autofluorescence in wt (630Δ*erm*), Δ*sigB* and *spo0A*::CT mutants. Dot plots of mean longitudinal green fluorescence (a.u.) of the analysed *C. difficile* 630Δ*erm* cells (green), Δ*sigB* (orange) and *spo0A*::CT (blue), previously presented in Fig. 2 and 3. Black lines represent the median values. Quantifications were performed using MicrobeJ from at least two biological replicates for each condition.

